# The effect of host admixture on wild house mouse gut microbiota is weak when accounting for spatial autocorrelation

**DOI:** 10.1101/2023.05.26.542413

**Authors:** Dagmar Čížková, Lucie Schmiedová, Martin Kváč, Bohumil Sak, Miloš Macholán, Jaroslav Piálek, Jakub Kreisinger

## Abstract

The question of how interactions between the gut microbiome and vertebrate hosts contribute to host adaptation and speciation is one of the major problems in current evolutionary research. Using bacteriome and mycobiome metabarcoding, we examined how these two components of the gut microbiota vary with the degree of host admixture in secondary contact between two house mouse subspecies (*Mus musculus musculus* and *M. m. domesticus*). We used a large dataset collected at two replicates of the hybrid zone and model-based statistical analyses to ensure the robustness of our results. Assuming that the microbiota of wild hosts suffers from spatial autocorrelation, we directly compared the results of statistical models that were spatially naive with those that accounted for spatial autocorrelation. We showed that neglecting spatial autocorrelation can drastically affect the results and lead to misleading conclusions. The spatial analyses showed little difference between subspecies, both in microbiome composition and in individual bacterial lineages. Similarly, the degree of admixture had minimal effects on the gut bacteriome and mycobiome and was caused by changes in a few microbial lineages that correspond to the common symbionts of free-living house mice. In contrast to previous studies, these data do not support the hypothesis that the microbiota plays an important role in host reproductive isolation in this particular model system.

## Introduction

Animal bodies are inhabited by diverse microbial communities that massively influence various aspects of host biology (Margulis, 1990). Understanding how host-microbiota interactions contribute to host adaptation, the emergence of new species, and the maintenance of species integrity and diversity is currently one of the most important problems in evolutionary and conservation biology (Sharpton, 2018; Zhu et al., 2021). An enormous research effort has been devoted to studying microbiota that inhabit the vertebrate digestive tract, where bacteria represent the major component, but other organisms such as fungi, protists, archaea, and viruses are also represented. Interactions between vertebrate hosts and gut bacteria are fundamental to both partners and are not limited to their metabolic interdependence, but also involve the host neuroendocrine and immune systems (McFall-Ngai et al., 2013). They are critical throughout the host’s life for proper development and long-term homeostasis of its physiology (Kundu et al., 2017). From an evolutionary perspective, these mutualistic interactions can occur within bacteria-host associations, spanning evolutionary time scales and linking the fitness interests of symbiotic partners (Groussin et al., 2020). This tight link between bacteria and their hosts is attributed to the part of the gut bacteriome that is transmitted long-term and stably across host generations (Shapira, 2016). There are numerous mechanisms that control and regulate the composition and function of the gut bacteriome, many of which are linked in some way. These include external factors, most notably dietary composition (Ley et al., 2008; Muegge et al., 2011), and host-internal factors, such as specific regulation by immune functions and more general regulation by the gut environment (Beasley et al., 2015; Sharp & Foster, 2022; Zheng et al., 2020). In addition, interactions between members of the microbial community also play an important role. Not surprisingly, disruption of these regulations can lead to dysbiosis of bacterial communities, which can trigger host dysfunction and thus reduce host fitness. If such disruption occurs as a direct result of admixture between divergent host populations, the reduction in fitness caused by dysbiosis of the microbiota may contribute to reproductive isolation of the two populations and ultimately promote their speciation (Brucker & Bordenstein, 2013; Chandler & Turelli, 2014).

Compared to gut bacteria, far less attention has been paid to non-bacterial gut communities. However, a growing number of studies focusing on intestinal fungi reveal their importance to the host. In some respects, host-mycobiome interactions resemble those between hosts and bacteria. For example, gut fungi play a direct role in host metabolism (Mims et al., 2021), they can interact with host immunity, and mycobiome dysbiosis has been linked to some complex human diseases (X. V. Li et al., 2019). At the same time, trans-generation transmission has been demonstrated for some fungal species (Bliss et al., 2008). Interactions between gut bacteria and fungi have been frequently reported (Peleg et al., 2010), but only recently has it been shown that colonization by gut fungi can causally influence the gut bacteriome and associated host phenotypes (van Tilburg Bernardes et al., 2020). On the other hand, the temporal stability of individual mycobiomes is low compared to the bacteriome, and their interindividual variability is high (Nash et al., 2017; Sharma et al., 2022; Wampach et al., 2017), likely due to stronger environmental influences, particularly host diet (David et al., 2014; Hallen-Adams et al., 2015; Hoffmann et al., 2013). Although diet composition can alter resident gut fungi in a substrate-dependent manner, it is suggested that fungi that are ingested in the diet but not integrated into microbial communities may cause the observed pattern and mask the signal from autochthonous mycoboime (Auchtung et al., 2018; Suhr & Hallen-Adams, 2015).

Most of our knowledge of host-microbiota interactions comes from experiments with captive animals or correlative studies on humans from industrialized countries. However in captive animal models, the microbiota exhibits significant changes compared to their free-living counterparts, resulting in altered microbiota-mediated functions and loss of certain beneficial traits under captive conditions (McKenzie et al., 2017; Rosshart et al., 2017). Similarly, the human microbiota has undergone massive changes following the transition to a Western lifestyle in industrialized countries, resulting in the depletion of some microbial species that are highly abundant in populations with traditional lifestyles (Gomez et al., 2016, 2019). These changes are thought to be related to a disruption of host-microbiota interactions that have evolved over long periods of close mutual coexistence, calling into question the suitability of these systems for evolutionary studies. Clearly, extensive empirical data from free-living populations are urgently needed, but obtaining and analysing such data presents a number of challenges. One of the most common and important causes of the complexity of these data is the fact that free-living individuals are typically sampled from spatially separated locations with varying geographic proximity. It has been regularly observed that samples from nearby locations have more similar microbiota than samples from geographically distant areas (Linnenbrink et al., 2013; Moeller et al., 2013, 2017; Rothschild et al., 2018) - a phenomenon also known as spatial autocorrelation. Spatial autocorrelation of the microbiota often occurs when some biotic (e.g., food resources, syntopic host communities) or abiotic factors (e.g., temperature, humidity) that shape microbial host communities lose similarity with increasing geographic distance. An important but often ignored consequence of this pattern is that microbiota samples derived from spatially proximate locations cannot be considered independent in a statistical sense. If spatial autocorrelation is not accounted for in statistical analyses, parameter estimates of statistical models may be biased and/or uncertainties around these parameters may be both inflated and reduced. In other words, failure to account for spatial autocorrelation can increase both Type I and Type II statistical errors and consequently lead to misleading conclusions. Unfortunately, there is a paucity of empirical studies examining the impact of spatial autocorrelation on analytical results in the context of host-associated microbiota research.

In this study, we investigated the consequences of neglecting spatial autocorrelation by analysing spatially organized gut microbiota data from two house mouse subspecies, *Mus musculus musculus* and *Mus musculus domesticus*, in a zone of their secondary contact. The two subspecies, which have diverged in allopatry for about 500 000 years (Y. Li et al., 2021; Phifer-Rixey et al., 2020), meet in Europe and form a ∼10 km narrow hybrid zone extending 2 500 km from Norway to the Black Sea (Ďureje et al., 2012; Jones et al., 2010). The zone is spatially structured with respect to the degree of mouse admixture - genetically pure parental subspecies at the edges with increasing degrees of genome admixture toward the center. This structure is relatively stable over time and is maintained by a balance between dispersal of admixing parents toward the center and selection against hybrids. Selection against hybrids is thought to be endogenous due to genetic incompatibilities, as evidenced by both genetic (Janousek et al., 2012; Macholán et al., 2007) and phenotypic studies (Albrechtová et al., 2012; Turner & Harr, 2014).

The first studies to address the role of mouse symbionts in the reproductive barrier reported that hybrid mice were more heavily parasitised by intestinal helminths compared to parental subspecies (Moulia et al., 1991; Sage et al., 1986), suggesting a possible role of host-symbiont interactions in the reduced fitness of hybrids. However, a larger scale model-based analysis of helminths in two replicates of the hybrid zone revealed an opposite pattern (Baird et al., 2012; Balard et al., 2020). These studies challenged the paradigm that the immune system is involved in incompatibilities that accelerate speciation. To date, only one study has attempted to investigate a possible role of the mouse gut microbiota in the reproductive isolation of house mice in the hybrid zone (J. Wang et al., 2015). This study showed that the gut bacterial communities of hybrid mice differ from the communities of parental mice. At the same time, this difference was correlated with changes in mouse immunophenotype and gut histopathology.

The main motivation of this study was to investigate how the gut microbiota of the house mouse varies with host genetic background (mouse subspecies and degree of admixture) in the house mouse hybrid zone. We also examined the effects of other host traits not related to host evolutionary divergence (sex, gravidity, and body weight). We used an extensive dataset collected at two different replicates of the hybrid zone (Czech and Regensburg transects), which allowed us to examine the effects of geography on microbiota variation at a larger scale and, importantly, to reduce the risk of confounding locally specific variation with systematic effects. We also compared patterns of microbiota variation between the two major components of gut microbial communities-the relatively well-described gut bacteriome and the understudied gut mycobiome. Finally, rigorous statistical modelling allowed a direct comparison between the results of spatially explicit and spatially naive analyses. In this way, we were able to determine the consequences of neglecting spatial autocorrelation in this model system, which is also important for other free-living host populations and species.

## Materials and Methods

### Sample collection

Fecal samples from 471 wild house mice were collected in two house mouse hybrid zone replicates: the Czech transect and the Regensburg transect (Figure 1). Replicates included allopatric localities with genetically pure *Mus musculus musculus* and *M. m. domesticus* as well as localities with varying degrees of admixture. Mice were live-trapped in farms or in their immediate vicinity and individually caged. After a maximum of 24 hours, they were anesthetized and dissected in a field laboratory using sterile tools. Kidneys and muscles were frozen in liquid nitrogen for allozyme typing, spleens were preserved in ethanol for genotyping, and colons, including feces, were preserved in ethanol for mycobiome and bacteriome metabarcoding. Mouse sex, body weight, and female gravidity were recorded during the dissections.

**Figure 1:**
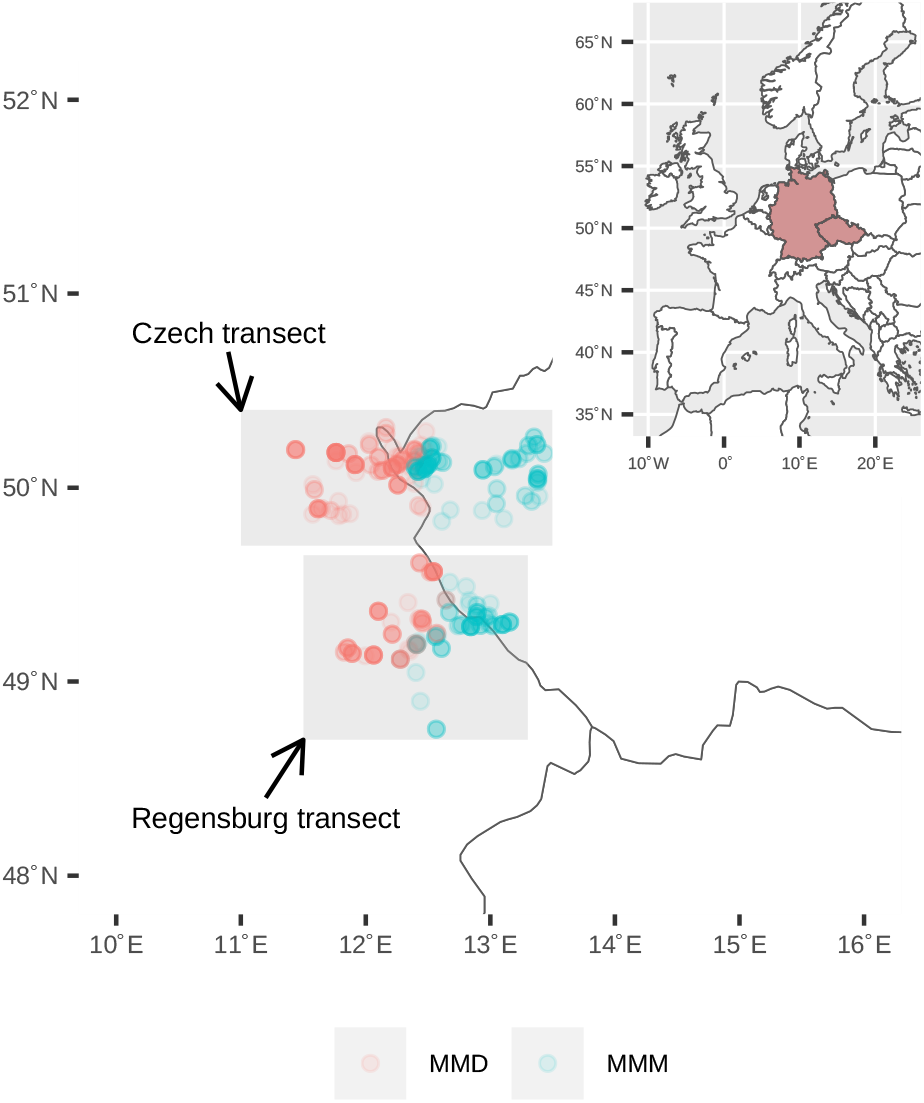
Map of sampling. The map shows two replicates of the hybrid zone, the Czech transect and the Regensburg transect. The circles represent the sampling sites, and the colour transparency is inverse to the number of individuals sampled. The colours represent the subspecies of the house mouse: *M. m. domesticus* (red; MMD) and *M. m. musculus* (blue; MMM).

### Mouse genotyping

The genetic background of each mouse was estimated as the proportion of *M. m. musculus* alleles (i.e., hybrid index), using two sets of diagnostic markers. The first set included 14 genetic and allozyme markers (Božíková et al., 2005; Dufková et al., 2011; Ďureje et al., 2012; Macholán et al., 2007; Munclinger et al., 2002, 2003). The second set consisted of more than 1400 X-linked and autosomal SNPs (L. Wang et al., 2011). While all individuals were genotyped with the first set of markers, SNPs were genotyped in a subset of individuals (n=271). The tight correlation between the individual-level hybrid index calculated with the two sets of markers (Pearson correlation: r = 0.980, d.f. = 270, p < 0.0001) suggested negligible variation between the two genotyping methods. Therefore, the average values of the hybrid index for individuals genotyped by both methods were considered. The hybrid indices, which ranged from 0 (pure *domesticus*) to 1 (pure *musculus*), were converted to the degree of admixture, which ranged from 0 (pure *musculus* and *domesticus*) to 0.5 (equal proportion of both genomes). The subspecies of each individual was determined based on the dominant genetic background.

### Microbiome metabarcoding

Fecal DNA was extracted using the QIAamp DNA Stool Mini Kit (QIAGEN) at the Institute of Parasitology, Biology Centre, CAS, as described elsewhere (Sak et al., 2011). As we have recently shown, the microbiota of house mouse feces is closely correlated with the microbiota of the lower intestinal tract and represents a suitable proxy for communities in the cecum and colon (Čížková et al., 2021). Gut bacteriome and mycobiome metabarcoding libraries were prepared using two-step PCR protocol at the Studenec Research Facility of the Institute of Vertebrate Biology, CAS, as described in Bendová et al. (2020) and Čížková et al. (2021). Briefly, standard metabarcoding primers of Klindworth et al. (2013) for bacterial 16S rRNA and ITS3 /ITS4 primers of White et al. (1990) flanking fungal ITS2 were used in the first PCR to amplify the specific rRNA loci. Dual indexes were introduced in t5he second PCR (in combination with inline barcodes, they allowed unique identification of each sample), and Illumina-compatible Nextera-like sequencing adapters were reconstituted. Each metabarcoding PCR was performed in a technical duplicate to account for PCR and sequencing stochasticity. PCRs in which amplification products were visible at both duplicates on the electrophoresis gel were pooled according to their concentration. In some fungal PCRs, amplification yield was low (whereas bacterial PCRs from the same samples amplified normally), likely due to the low amount of fungal DNA in these samples. Therefore, greater variation in the molar fractions of pooled PCRs for fungi was expected. Pools were sequenced using Illumina Miseq (v3 kit, 300 bp paired-end reads) at CEITEC, Brno, Czech Republic.

### Analyses of microbiome diversity and composition

Skewer (Jiang et al., 2014) was used to demultiplex samples and detect and then trim gene-specific primers. Low-quality reads (expected error rate per paired-end read > 2) were eliminated, quality-filtered reads were denoised, and abundance matrices were generated using the read counts for each 16S rRNA and ITS2 amplicon sequencing variant (hereafter ASV) in each sample using the R package Dada2 (Callahan et al., 2016). Subsequently, uchime (Edgar et al., 2011) was used to detect and eliminate chimeric ASVs. The gold.fna (available at: https://drive5.com/uchime/gold.fa) and UNITE v. 02.02.2019 (Kõljalg et al., 2013) databases were used as references for filtering chimeras from the gut bacteriome and mycobiome datasets, respectively. After eliminating chimeric ASVs, the taxonomy for non-chimeric ASVs was assigned with 80% posterior confidence by the RDP classifier (Q. Wang et al., 2007). The Silva database v.138, (Quast et al., 2013) was used for bacterial ASV annotations and the UNITE database for fungal ASV annotations. Using Procrustean analysis, we checked the consistency of the composition of ASV profiles between technical duplicates and excluded samples where the duplicates corresponding to a given sample showed excessive compositional divergence. We then merged the duplicated data while eliminating all ASVs that were not detected in both duplicates. Finally, we excluded all samples with a number of high-quality reads < 1000. A total of 7 samples were removed for the bacteriome and 33 samples for the mycobiome, resulting in 464 and 438 samples in the final dataset for bacteriome and mycobiome, respectively. For each sample, we calculated microbial alpha diversity using the Shannon diversity index and the number of observed ASVs using rarefied ASVs abundance matrices (rarefaction threshold = 1038 for bacteriome and 1085 for mycobiome dataset; i.e., the minimum sequencing depth reached). To quantify dissimilarities in the composition of microbial communities, two types of beta diversity indexes were calculated: Bray-Curtis index based on relative ASVs abundances and Jaccard index which accounts for ASV presence/absence only.

### Statistical analyses

Statistical analyses were performed to evaluate how different host-associated variables explain microbiome variation, whether the resulting effects differ between spatially aware and spatially naive models, and whether models incorporating spatial information fit our data better compared with non-spatial models. The microbiome variables included as response variables in the analyses were: alpha diversity of microbial communities (i.e., Shannon diversity index and the number of observed ASVs), beta diversity of microbial communities (i.e., differences in microbiome composition expressed as Jaccard and Bray-Curtis distances), and prevalence (i.e., presence/absence) or relative abundance of individual microbial ASVs. Host-associated variables considered predictors of microbiota variation included mouse subspecies (*musculus* or *domesticus*), degree of admixture (continuous variable, ranging from 0 for both pure subspecies to 0.5 for equally mixed genomes), hybrid zone replicate (Czech or Regensburg transect), body weight (continuous variable), sex, and gravidity (factorial predictor with three levels: male, female gravid, and female non-gravid). In addition, the interaction between subspecies and degree of admixture was investigated. Sampling location was included as a random effect in both the spatially naive and spatially aware analyses. However, in the spatially aware models that accounted for spatial autocorrelation in microbiome variation, geographic distances between sampling sites were also considered. All of these analyses, detailed below, were performed separately for bacterial and fungal microbial communities.

#### a) Analyses of microbial alpha diversity

Shannon diversity and the number of observed ASVs were included as response variables in mixed models from the R package spaMM (Rousset & Ferdy, 2014). The number of observed ASVs was log_10_-transformed to achieve the Gaussian distribution of residuals. Model predictors are explained above. In spatially explicit version of these models, the spatial autocorrelation was modelled using the Matern correlation structure for geographic coordinates. By stepwise elimination of non-significant terms a minimal adequate model was built. Significance testing of predictors was based on likelihood ratio tests. Fit of the two model versions - spatial and non-spatial - were compared using the conditional Akaike information criterion (AIC) estimated by the bootstrap method (Saefken et al., 2014).

#### b) Analyses of microbial beta diversity

We employed two parallel approaches for these analyses. In the first "major gradients approach," we performed principal coordinate analysis (PCoA) for the Jaccard and Bray-Curtis dissimilarities and used the scores for the first two PCoA axes as response variables in mixed models, which had the same structure as described for alpha-diversity.

In the second "whole community" approach, the Jaccard and Bray-Curtis indexes were used as response variables in Multivariate Distance Matrix Regression mixed model (R package MDMR; (McArtor et al., 2017). In spatially aware models, geographic distances between sites scaled by the principal coordinates of the neighbourhood matrix (PCNM) were used as spatial covariates. To prevent MDMR overfitting, only the PCNM gradients exhibiting significant correlation with Jaccard or Bray-Curtis indexed were included. These gradients were identified by distance-based redundancy analyses (db-RDA) constructed using a stepwise forward selection procedure using the ordistep function in the vegan R package (Oksanen, 2010).

#### c) Analyses of microbial ASVs

Joint species distribution models (JSDM) from the R package HMSC (Tikhonov et al., 2020) were used for these analyses. JSDMs assumed random variation among sites and samples. In addition, the spatially aware models included also geographic distances. We applied the hurdle modelling approach, i.e., we first analysed variation in the prevalence (i.e. presence/absence) of each ASV assuming a binary distribution and then focused on variation in the abundance of each ASV in the subset of the samples where its presence was detected, assuming lognormal Poisson distribution. JSDMs were run on two Markov Chain Monte Carlo (MCMC) chains with 5 500 000 iterations each. The first 500 000 iterations were discarded as burning and the thinning intervals were fixed at 5 000 steps. Convergence of the chains for each parameter was assessed by visual inspection of the trace plots and by the corresponding effective sample sizes and Gelman-Rubin convergence diagnostics. Support for the estimated parameters was assessed using 95% credible posterior intervals.

## Results

### Microbial alpha diversity

After all filtering steps, the gut bacteriome dataset contained 464 samples and 8 689 bacterial ASVs (average ASV richness = 69.5 [SD = 28.57198]) represented by 3 034 185 reads (average sequencing depth = 6 539 sequences per sample, range = 1 038 - 21 072). Mixed models for alpha diversity that accounted for spatial autocorrelation received very similar support compared with those that did not account for the effect of spatial proximity and only accounted for variation between sampling sites via the random effect (AIC = −201.8533 and −201.8533 for spatial vs. non-spatial models for ASVs richness; AIC = 791.7882 and 791.0379 for spatial vs. non-spatial models for Shannon diversity). Consequently, the parameter estimates for the predictor variables and their significances were very similar for both model versions (Table 1, S1). Both ASVs richness and Shannon diversity were significantly increased in the Czech transect compared to the Regensburg transect (Table S1), whereas the other alpha diversity predictors were not significant (Table 1).

**Table 1:** Variation in microbial alpha diversity. The full model (including all predictors) was reduced by stepwise backward elimination of nonsignificant predictors to obtain a minimal adequate model containing only significant predictors (boldface). Significance tests were based on likelihood ratio tests, in which significance (p) was derived from the deviance changes between models (χ^2^) and the corresponding degrees of freedom (Δ D.f.). Variation in two alpha diversity measures (Shannon index and Observed ASVs) was assessed separately for bacterial and fungal communities. Models with and without spatial autocorrelation were evaluated in parallel.

| Community | Response | Predictor | $\Delta$ D.f. | Spatial models | | Non-spatial models | |
| --- | --- | --- | --- | --- | --- | --- | --- |
| | | | | $\chi^2$ | p | $\chi^2$ | p |
| Bacteriome | Observed | Subspecies | 1 | 0,9952 | 0,3185 | 0,9952 | 0,3185 |
|  |  | Admixture | 1 | 0,0504 | 0,8224 | 0,0504 | 0,8224 |
|  |  | Subspec x Admix | 1 | 0,0570 | 0,8113 | 0,0570 | 0,8113 |
|  |  | Body weight | 1 | 0,8139 | 0,3670 | 0,7181 | 0,3968 |
|  |  | Sex+Gravidity | 2 | 1,7230 | 0,4225 | 1,8188 | 0,4028 |
|  |  | Zone replicate | 1 | **7,6013** | **0,0058** | **15,6292** | **0,0001** |
|  | Shannon | Subspecies | 1 | 0,5369 | 0,4637 | 0,5384 | 0,4631 |
|  |  | Admixture | 1 | 2,3829 | 0,1227 | 2,3829 | 0,1227 |
|  |  | Subspec x Admix | 1 | 0,4626 | 0,4964 | 0,4626 | 0,4964 |
|  |  | Body weight | 1 | 0,0600 | 0,8066 | 0,0600 | 0,8065 |
|  |  | Sex+Gravidity | 2 | 0,2177 | 0,8969 | 0,2177 | 0,8968 |
|  |  | Zone replicate | 1 | **4,2863** | **0,0384** | **4,2850** | **0,0385** |
| Mycobiome | Observed | Subspecies | 1 | 1,7296 | 0,1885 | 1,9047 | 0,1676 |
|  |  | Admixture | 1 | 0,0648 | 0,7990 | 0,0556 | 0,8136 |
|  |  | Admix x Subspec | 1 | 0,0088 | 0,9253 | 0,0073 | 0,9318 |
|  |  | Body weight | 1 | 0,5648 | 0,4523 | 0,6969 | 0,4038 |
|  |  | Sex+Gravidity | 2 | 4,5912 | 0,1007 | 4,0433 | 0,1324 |
|  |  | Zone replicate | 1 | **6,7284** | **0,0095** | **16,5267** | **0,0000** |
|  | Shannon | Subspecies | 1 | **3,9337** | **0,0473** | **5,6779** | **0,0172** |
|  |  | Admixture | 1 | 1,0345 | 0,3091 | 1,4951 | 0,2214 |
|  |  | Admix x Subspec | 1 | 1,3929 | 0,2379 | 1,1870 | 0,2759 |
|  |  | Body weight | 1 | **7,4355** | **0,0064** | **7,4355** | **0,0064** |
|  |  | Sex+Gravidity | 2 | 2,4176 | 0,2986 | 2,4621 | 0,2920 |
|  |  | Zone replicate | 1 | 0,0034 | 0,9532 | 0,3420 | 0,5587 |

For gut mycobiome we detected a total of 1 140 fungal ASVs in 438 gut samples, with an average ASV richness per sample of 13.1 (s.d. = 7.9) and an average sequencing depth of 8 230 reads per sample (range = 1 085 - 51 573, total number of high-quality reads = 3 604 543). In contrast to the bacterial alpha diversity analyses, spatially explicit models for fungal alpha diversity received higher support compared to models that did not account for spatial autocorrelation (AIC = 182.4014 vs. 186.2871 for spatial vs. non-spatial models for ASVs richness; AIC = 1005.1278 vs. 1009.1008 for spatial vs. non-spatial models for Shannon diversity).

In both spatial and non-spatial models, fungal ASV richness was increased in the Czech transect, and furthermore, Shannon alpha diversity was higher in *M. m. musculus* and decreased with increasing body mass. However, the non-spatial models showed inflated estimates for the effect of transects and mouse subspecies, i.e. predictors showing a clear geographic pattern of variation (Table 1, S1).

### Gut bacteriome beta diversity

Bacteriome taxonomic composition at the class (Figure 2) and genus (Figure S1) levels showed no pronounced differences between the two house mouse subspecies, the two hybrid zone replicates, or due to genetic admixture. PCoA ordination (Figure 3) also indicated little variation in gut bacteriome composition at the ASV level between transects or subspecies.

**Figure 2:**
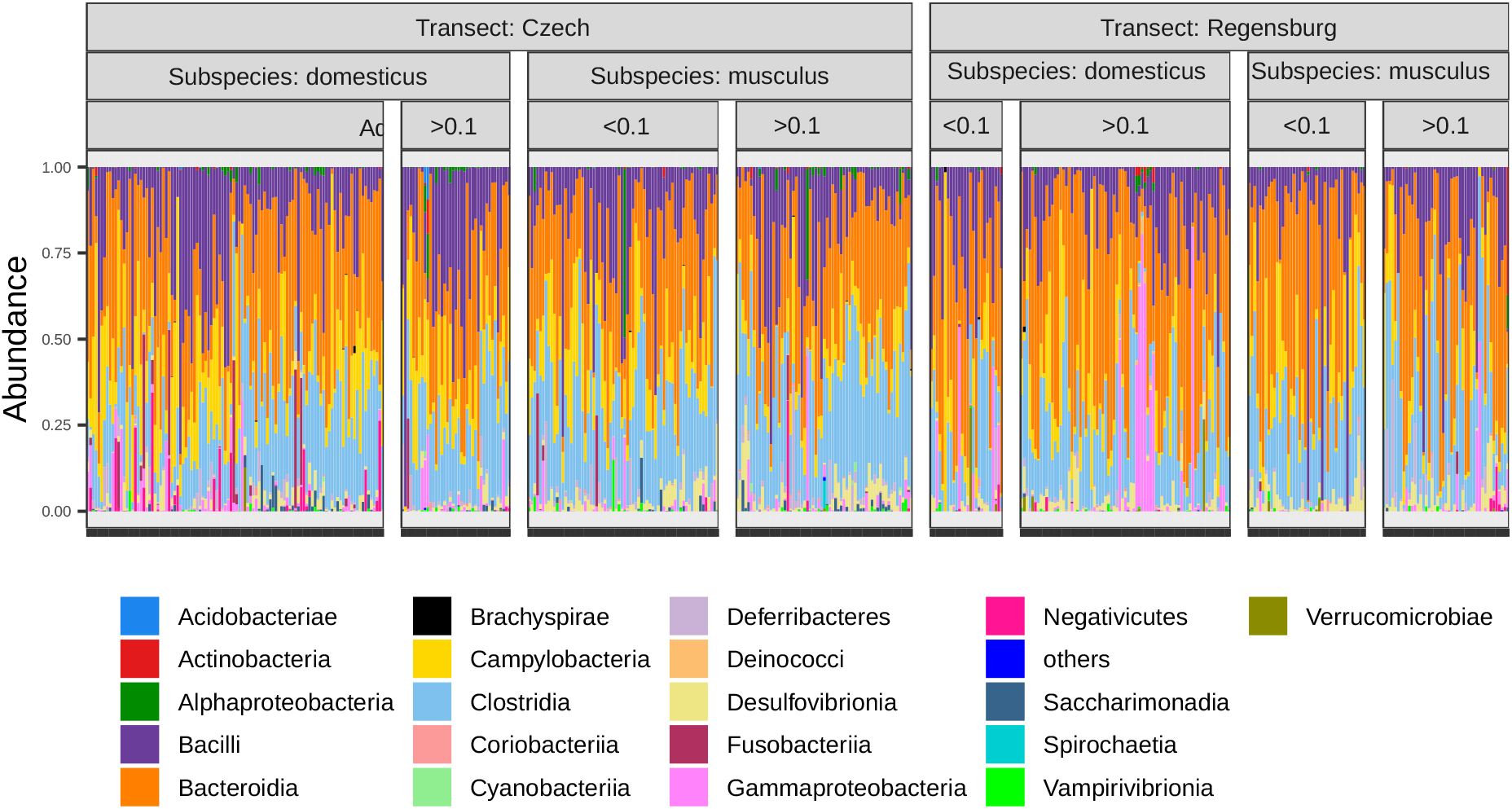
Composition of gut bacteriome. Proportion of bacterial classes detected in the gut microbiota of house mice. Profiles were sorted by hybrid zone replicate (Transect: Czech or Regensburg), house mouse subspecies (Subspecies: musculus or domesticus), and degree of admixture (Admixture < 0.1 or > 0.1; admixture was divided into the two categories for display purposes only, otherwise it was used as a continuous variable).

**Figure 3:**
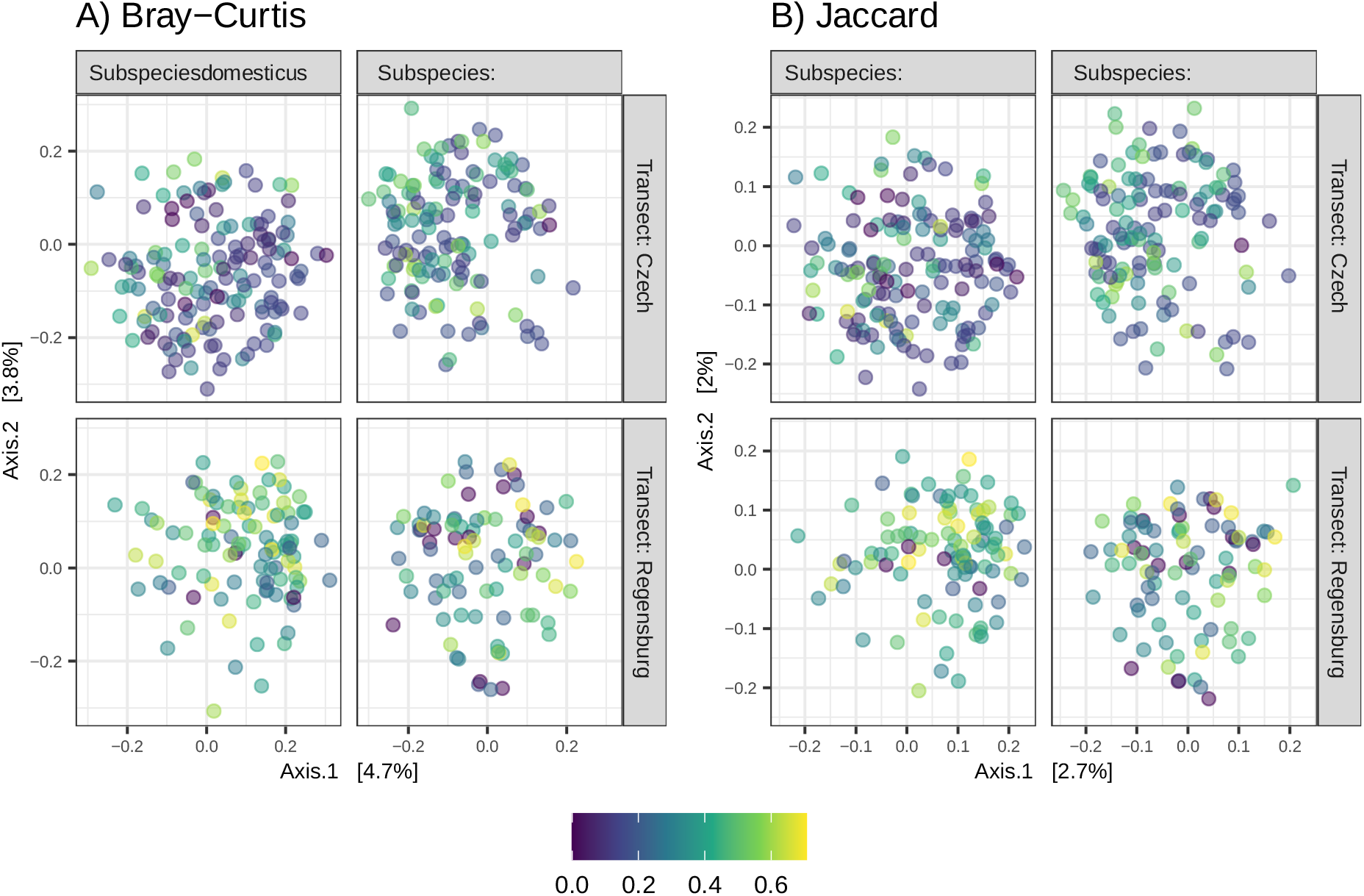
Ordination of gut bacteriome samples. PCoA ordination shows divergence in gut bacteriome composition between replicates of the hybrid zone (Transect: Czech and Regensburg), subspecies of house mouse (Subspecies: musculus and domesticus) and due to genetic admixture (colour scale expresses degree of admixture after square root transformation, suppressing influence of extreme values). Ordination was performed for dissimilarities between samples considering A) variation in relative ASV frequencies (Bray-Curtis) and B) presence/absence of ASVs (Jaccard).

The results of the "major gradients" analyses showed that mixed models for the scores of the first two PCoA axes that accounted for spatial autocorrelation had a better fit (i.e., lower AIC values) compared with their non-spatial counterparts (Table S3). According to the models with spatial autocorrelation, bacteriome composition varied with transects and subspecies. In addition, a significant correlation was observed between body weight and scores for the first PCoA axis. Finally, models for the second PCoA axis and Bray-Curtis dissimilarities revealed significant interactions between the degree of admixture and subspecies, suggesting that the microbiota of *musculus* and *domesticus* mice may exhibit subspecies-specific changes in admixed individuals (Table S2, S4). We next examined in more detail whether and how the inclusion of spatial autocorrelation affected the results of the fitted models. We found no systematic bias in the absolute values of parameter estimates (paired t-test: t = 0.8875, df = 27, p = 0.3827) in models with or without spatial autocorrelation. However, predictors in models without spatial autocorrelation were associated with significantly higher likelihood ratio values (paired t-test: t = −2.3100, df = 23, p = 0.0302) and their estimates had lower uncertainty (paired t-test: t =2.7380, df = 27, p = 0.0108), indicating anti-conservative significance values. This was especially true for predictors that were closely related to the spatial distribution of samples, i.e., hybrid zone transect, subspecies, and in some cases also degree of admixture.

Because the first two PCoA axes explained only a small portion of the total variation in gut bacteriome composition, we also applied the "whole community" approach. According to the db-RDA, spatial proximity was found to be an important factor influencing gut bacteriome composition (F_(14,119)_ = 1.6162, p < 0.001, adjusted R^2^ = 0.060 for Bray-Curtis and F_(12,121)_ = 1.3864, p < 0.001, adjusted R^2^ = 0.0330 and for Jaccard dissimilarities), confirming the relevance of spatially aware models. Consistent with the "major gradients" results, MDMRs that controlled for spatial effect yielded lower significance for most predictors than spatially naive MDMRs (paired t-test: t = - 2.8945, df = 11, p = 0.0146). Of note, the effect of admixture, which was highly significant based on spatially naive MDMRs, was far from significant when PCNM axes were included as covariates, and the same was true for the admixture x subspecies interaction (Table 2).

**Table 2:** Variation in microbiota composition: whole community approach. Bray-Curtis and Jaccard dissimilarities were used as response variables for mixed MDMR models in which variation between sample locations was modeled as a random effect. Models that included spatial dissimilarities between sample locations (i.e., scores for PCNM axes) as covariates were fit in parallel to models without spatial effects. Values of significant predictors are in bold.

| Community | Response | Predictor | $\Delta$ D.f. | Spatial models | | Non-spatial models | |
| --- | --- | --- | --- | --- | --- | --- | --- |
|  |  |  |  | Test stat. | p | Test stat. | p |
| Bacteriome | Bray-Curtis | (Intercept) | 1 | 1,9233 | 0,0002 | 1,9055 | 0,0002 |
|  |  | Subspecies | 1 | **1,7436** | **0,0010** | **3,4523** | **0,0000** |
|  |  | Admixture | 1 | 1,1875 | 0,1420 | **1,7513** | **0,0009** |
|  |  | Subspec x Admix | 1 | 1,3028 | 0,0576 | **1,4295** | **0,0193** |
|  |  | Body weight | 1 | **2,1305** | **0,0000** | **2,1847** | **0,0000** |
|  |  | Sex+Gravidity | 2 | 1,9514 | 0,5480 | 1,9354 | 0,5743 |
|  |  | Zone replicate | 1 | **1,9328** | **0,0001** | **2,5899** | **0,0000** |
|  | Jaccard | (Intercept) | 1 | 1,2630 | 0,0093 | 1,4152 | 0,0005 |
|  |  | Subspecies | 1 | 1,1521 | 0,0667 | **2,1424** | **0,0000** |
|  |  | Admixture | 1 | 1,0750 | 0,2077 | **1,3462** | **0,0018** |
|  |  | Subspec x Admix | 1 | 1,0622 | 0,2443 | **1,2118** | **0,0239** |
|  |  | Body weight | 1 | **1,4769** | **0,0001** | **1,4883** | **0,0001** |
|  |  | Sex+Gravidity | 2 | 2,0529 | 0,3331 | 2,0436 | 0,3575 |
|  |  | Zone replicate | 1 | **1,3773** | **0,0010** | **2,1262** | **0,0000** |
| Mycobiome | Bray-Curtis | (Intercept) | 1 | 1,3667 | 0,1309 | 1,8440 | 0,0235 |
|  |  | Subspecies | 1 | 1,1006 | 0,3162 | **4,0746** | **0,0000** |
|  |  | Admixture | 1 | 1,3743 | 0,1275 | **2,1071** | **0,0090** |
|  |  | Subspec x Admix | 1 | 1,0589 | 0,3589 | **2,3084** | **0,0043** |
|  |  | Body weight | 1 | **2,0269** | **0,0120** | **2,0219** | **0,0122** |
|  |  | Sex+Gravidity | 2 | 1,3674 | 0,9374 | 1,4527 | 0,8963 |
|  |  | Zone replicate | 1 | **3,9332** | **0,0000** | **12,3854** | **0,0000** |
|  | Jaccard | (Intercept) | 1 | 1,2036 | 0,1199 | 1,2070 | 0,1168 |
|  |  | Subspecies | 1 | 1,0835 | 0,2801 | **2,2154** | **0,0000** |
|  |  | Admixture | 1 | **1,4396** | **0,0163** | **2,1210** | **0,0000** |
|  |  | Subspec x Admix | 1 | 1,0471 | 0,3501 | **1,5693** | **0,0049** |
|  |  | Body weight | 1 | 1,0659 | 0,3128 | 0,9831 | 0,4946 |
|  |  | Sex+Gravidity | 2 | 1,8723 | 0,6811 | 1,8784 | 0,6710 |
|  |  | Zone replicate | 1 | **5,5277** | **0,0000** | **13,0785** | **0,0000** |

### Gut mycobiome beta diversity

According to the taxonomic composition (Figure S1), *Kazachstania* represented the dominant fraction of the mouse mycobiome with an average percentage of 32.7%. We also detected relatively high abundances of unassigned Ascomycota (average 14.1% of reads), *Aspergilus* (6.5%), *Candida* (6.5%), and *Penicillium* (5.7%). A significant effect of geographic distances between sampling sites on gut mycobiome composition was detected by db-RDA (F_(15,113)_ = 2.6305, p < 0.001, adjusted R^2^ = 0.1561 for Bray-Curtis and F_(14,114)_ = 1.8255, p < 0.001, adjusted R^2^ = 0.0763 and for Jaccard dissimilarities).

Spatial autocorrelation in fungal composition was also detected by the "major gradients" analyses testing gut mycobiome variation along PCoA gradients and was implied by higher support for spatially explicit mixed models compared with models that ignored information about spatial proximity of samples (Table S3). Nonetheless, both spatial and non-spatial model versions typically identified a similar set of predictors significantly affecting fungal community composition (Table 2, S2), e.g., the main effect of admixture or its interaction with subspecies identity, along with the effect of transect or gravidity. A notable discrepancy was observed for the second PCoA axis, where the spatial model detected a significant effect of body weight, whereas the non-spatial model suggested a significance of mouse subspecies. Parameter estimates from the spatially explicit models were associated with greater uncertainty (paired t-test: t = 3.0644, df = 27, p-value = 0.0049), but this did not result in a decrease in test statistics during the model selection process (t = 0.1682, df = 23, p-value = 0.8679). Also, no systematic bias in absolute values of estimated parameters was observed between spatial and non-spatial models (t = −0.6577, df = 27, p-value = 0.5163) Predictors in spatial "whole community" analyses had lower test statistic values compared to spatially naive MDMRs (paired t-test: t = −2.2744, df = 11, p-value = 0.0440). As a result, the variation between the two subspecies or due to the interaction between subspecies and admixture, which was highly significant for both Bray-Curtis and Jaccard dissimilarity for the non-spatial MDMRs, was far from significant when the MDMRs were controlled for the effect of spatial proximity. On the other hand, variation between transects was supported by both spatial and non-spatial MDMRs, regardless of the dissimilarity measure. In addition, both spatial and non-spatial MDMRs for Jaccard dissimilarities indicated variation in composition due to the main effect of admixture and MDMRs for Bray-Curtis dissimilarities showed significant variation due to the effect of body weight (Table 2).

### ASV-level microbiota variation

The spatially explicit and non-spatial versions of the JSDM for the abundance of the individual bacterial ASVs generally yielded similar parameter sets with a high degree of posterior support (> 95%, Figure 4). According to these analyses, variation in the abundance of bacterial ASVs was mostly associated with differences between the hybrid zone replicates (eight ASVs differed between the two transects in spatial models) or body weight (four ASVs). The abundance of two ASVs varied between subspecies, but there was no support for the effect of admixture, sex, and/or gravidity. Compared with the abundance-based JSDM, the analyses that considered the prevalence (i.e. presence/absence) of bacterial ASVs revealed a higher number of highly supported correlations between the ASVs tested and the set of predictors. In addition, the prevalence based analyses showed greater differences between spatially naive and spatially explicit models compared to the abundance. For example, the prevalence of 14 bacterial ASVs varied between the mouse subspecies according to the non-spatial JSDM, whereas there were only seven ASVs in the spatial JSDM. Similarly, only six ASVs were affected by admixture or interaction between admixture and subspecies in the spatial JSDM, compared to eight ASVs in the case of the non-spatial JSDM. The prevalence of 16 and 15 ASVs varied between transects for the spatial and non-spatial models, respectively. We also found an effect of body mass, however, there was no variation associated with sex or gravidity (Figure 4).

**Figure 4:**
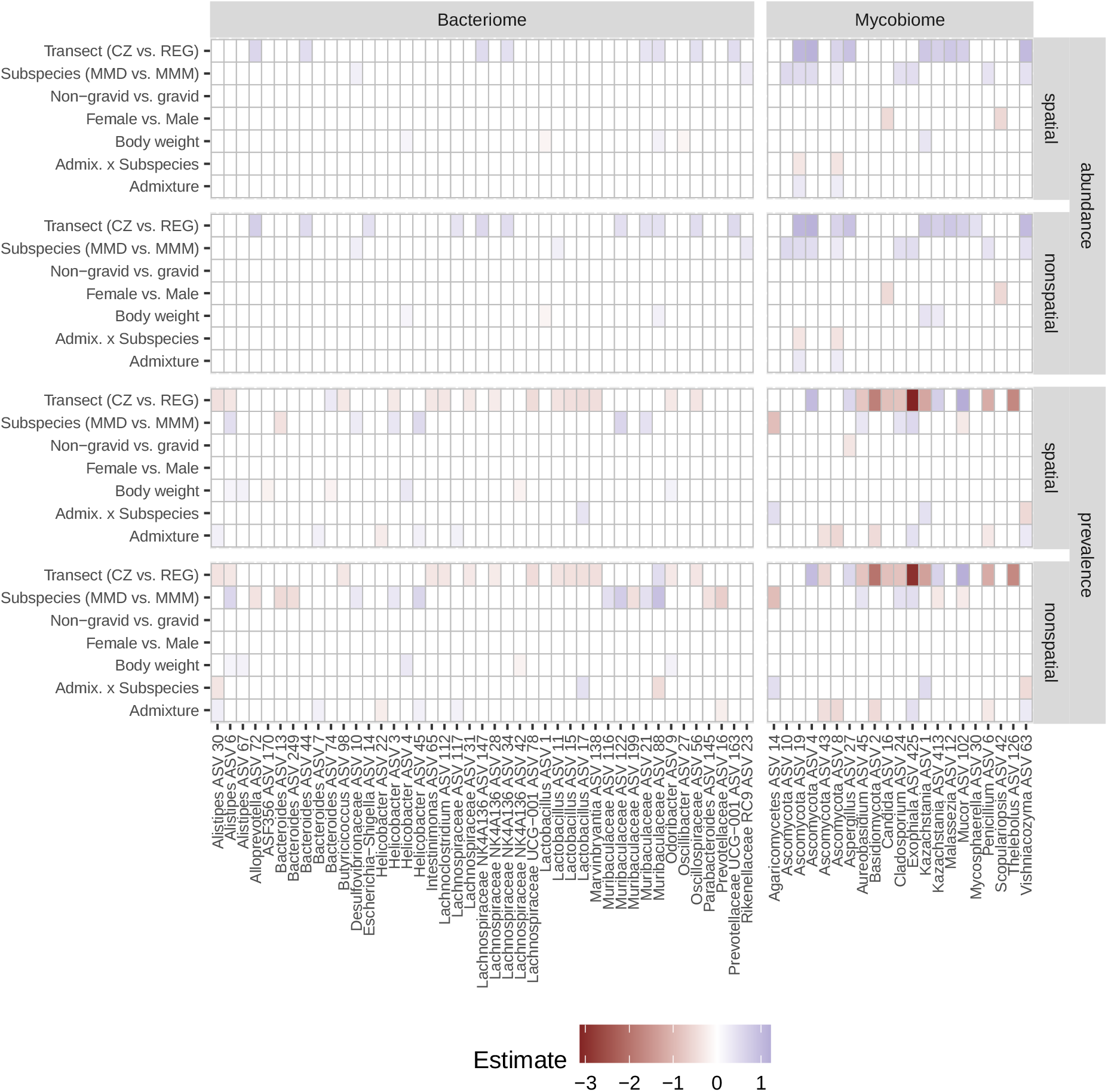
Results of joint species distribution models. Heatmap shows parameter estimates for model predictors (CZ = Czech transect, REG = Regensburg transect, MMM = M. m. musculus, MMD = M. m. domesticus) and individual bacterial or fungal ASVs. Separate HMSC analyses were performed with and without consideration of spatial autocorrelation (spatial and nonspatial). Variation in the prevalence or abundance of ASVs was modelled separately. Parameters that did not receive high posterior support (i.e., < 0.95) are indicated by white cells. Only ASVs in which posterior support was high for at least one predictor are shown.

Consistent with JSDM for the gut bacteriome, ASV-level variation in the fungal community was mainly influenced by differences between hybrid zone replicates (nine ASVs for abundance and 12 ASVs for prevalence, with spatial models) and mouse subspecies (eight ASVs for abundance and five ASVs for prevalence). In addition, several fungal ASVs varied with the degree of genetic admixture and body mass. Contrary to gut bacteriome, we detected sex-dependent differences in the abundance or prevalence of two fungal ASVs and differences associated with gravidity (one ASVs) (Figure 4).

Based on a direct comparison of spatial and non-spatial JSDM results, we found that for bacterial ASV abundance, all parameter estimates in non-spatial models were inflated (Table S5). In the case of JSDMs that considered prevalence of bacterial ASVs or abundance of fungal ASVs, spatially naive models also resulted in inflated estimates for some predictors directly related to sample geography, e.g., admixture, subspecies, or their interaction. The only exception was a larger difference between the two hybrid zone transects in the prevalence of bacterial ASV as determined by the spatial model. Uncertainty in parameter estimates, as assessed by the width of the 95% posterior credibility intervals, was increased in the spatial models, especially in the case of predictors related to the geographic origin of samples, e.g., subspecies, transect, degree of admixture, whereas the opposite was true in the case of the effect of gravidity, which is not directly related to geography (Table S5).

## Discussion

The house mouse hybrid zone is an excellent system for studying the role of the microbiota in host evolution under natural selection. A potential drawback of this system is the inherent spatial structure of the zone, in which the host genetic background shows clinal variation in space, such that animals sampled close to each other tend to have similar genetic backgrounds. Further structuring can be introduced by sampling multiple individuals at the same site. In this system, sampling sites are represented by spatially separated human settlements (e.g., farms). Within these localities, mice form genetically related social groups, and their dispersal between localities is limited (Klein & Bailey, 1971; Pocock et al., 2004). Under these circumstances, the effects of host genetic background on variation in gut microbiota may be confounded with the effects of other factors shaping gut microbiota that exhibit some spatial gradient. For example, changes in external microbial pools (in the environment or in other sympatric hosts) or food resources are neither obvious nor easily observable factors, and little is known about their spatial distribution. Therefore, in the first part of our study, we carefully examined the effects of spatial proximity (spatial autocorrelation) on variation in gut microbiota. Our approach was based on rigorous statistical modelling and a direct comparison of non-spatial models that ignored spatial autocorrelation and spatial models that accounted for spatial autocorrelation.

We have shown that the results of statistical models can vary in complex ways depending on which level of microbial diversity is being examined (i.e., alpha diversity vs. microbiome composition vs. ASV-level analyses) and between two domains of microbial symbionts (i.e., bacteria vs. fungi). In the case of microbial alpha diversity, we found little difference between spatial and non-spatial models for both the gut bacteriome and the mycobiome (Table 1). However, for microbiome composition, spatial models showed better fit and consequently there were substantial differences between the results of spatial and non-spatial models (Table 2, S2). These mainly affected the predictors directly related to the spatial distribution of the analysed samples and were associated with underestimation of uncertainties around the estimated parameters in the non-spatial analyses, rather than systematic biases in the parameter estimates (Table S3, S4). In particular, differentiation of the microbiota by hybrid zone transects was generally less supported by the spatial models. However, differences between models were most pronounced for variables related to the genetic background of the mice (subspecies, admixture, and their interaction), which often lost statistical support in the spatially explicit analyses. An illustrative example was the effect of the degree of host admixture on bacteriome composition (Table 2, S2). While admixture had a highly significant effect on both composition measures in the non-spatial version of both the "major gradients" and the "whole community" bacteriome analyses, the effects of admixture became nonsignificant in spatial models. In contrast, the effects of other variables (sex, pregnancy, body weight) were highly consistent between the spatial and non-spatial analyses. The only exception was "unmasking" of the body weight effect in the spatial "whole community" analyses of bacterial prevalence and fungal abundance.

At the level of individual ASVs, the effect of spatial autocorrelation was more pronounced in the bacteriome data than in the mycobiome. However, in contrast to the microbiome composition, ASV discrepancies between spatial and non-spatial models were mainly due to differences in parameter estimates rather than differences in their uncertainty. Similar to the microbiome composition, the difference was that bacterial ASVs associated with host genetic background were dropped from the spatial model results; for example, half of the ASVs with different prevalence between subspecies in the non-spatial model was missing from the spatial results (Figure 4). At the same time, additional bacterial ASVs associated with host body weight were identified by the spatial model.

Although distances between sampling sites explained only a relatively small proportion of the total variance in microbial profiles, our analyses clearly showed that it is not safe to ignore even such weak spatial effects. Conversely, neglecting spatial autocorrelation can dramatically affect the results of statistical models and potentially lead to erroneous biological conclusions. Particularly, confidence in the impact of spatially nonrandom host variables on the host-associated microbiome (i.e., host genetic background in our case) can be overestimated. On the contrary, spatially aware models can reveal the effects of spatially independent variables that are otherwise hidden in spatially naive analyses (e.g., body weight). Because spatial autocorrelation is a common aspect of animal-associated microbiota data collected in free-living populations, our contribution suggests that this potentially important aspect should be carefully investigated and accounted for in statistical calculations.

Host populations with separate evolutionary histories and distinct geographic distributions are expected to have specific microbiota (Groussin et al., 2020), which is also true for the two allopatric subspecies of the house mouse, *M. m. musculus* and *M. m. domesticus*. The microbes that live and evolve in the bodies of their hosts and that are continuously transmitted across host generations co-diverge with their hosts. In addition to selectively neutral divergence, microbes can also adaptively differentiate in response to changes between host lineages. This includes, for example, host divergence in the mechanisms regulating the microbiota or in its ecology. If selection translates into suppressed/increased growth of a microbial lineage, the shift in abundance will affect the entire microbial community and lead to divergence in community composition. On the other hand, a uniform selection pressure over the host lineages stabilizes microbial community composition and function. Convergent microbial communities between divergent hosts may be associated, for example, with specific ontogenetic stages, sex, or similar ecological conditions (Delsuc et al., 2014; Kreisinger et al., 2014; Sang et al., 2022; Zhang et al., 2021). In contrast to transgenerational transmission, the specificity of horizontally transmitted microbes incorporated into the host microbiota from external sources (e.g., environment, other syntopic hosts, food chains) is largely determined by their dispersal abilities/opportunities (Moeller et al., 2017) and their adaptation to the external environment. Therefore, environmentally transmitted microbes can only be specific to allopatric hosts if the barriers to their dispersal and/or external selection pressures coincide with the geographic distribution of the host.

We first examined changes in microbial communities and individual microbial lineages between the two house mouse subspecies and compared them to changes between the two geographically separated hybrid zone replicates. We did not find a dramatic effect of host subspecies on gut microbiota composition that would have been evident in the exploratory ordination analyses (Figure 3 and S2), but both the "whole community" and "major gradients" model-based approaches revealed some differences between the subspecies. The effect of subspecies, although rather weak, was still significant for all measures of bacteriome composition after spatial correction. However, in the case of the mycobiome, differences between subspecies were driven by the spatial proximity of sampled mice, as evidenced by a loss of significance in spatial models compared to non-spatial models. Transect differences that affected both bacterial and fungal community composition and diversity, detected with both spatial and non-spatial models, suggested an environmental role independent of subspecies. Furthermore, differentiation of the mycobiome among the hybrid zone replicates was probably the only apparent pattern in the ordination diagrams (Jaccard in Figure S2). Microbial diversity was generally greater in the Czech transect, perhaps reflecting the greater diversity of the landscape and human settlements in this area.

At the level of individual bacterial lineages (Figure 4, S3), ASVs that differed among subspecies belonged to the phyla Bacteroidota, Thermodesulfobacteriota and, Campylobacterota and represented gram-negative, non-sporulating anaerobes with low oxygen tolerance characterized by high levels of transgenerational transmission (Ferretti et al., 2018; Lagkouvardos et al., 2019; Linz et al., 2007). Although some members of these phyla also varied among hybrid zone replicates, the majority of transect-dependent ASVs belonged to the Bacillota phylum (classes Bacilli and Clostridia). These gram-positive bacteria have low host specificity (Moeller et al., 2016), consistent with their ability to survive in the environment in the form of spores (Galperin et al., 2012). Therefore, the differences among transects appear to be due to environmentally transmitted microbes.

Fungal ASVs that differed among subspecies included mainly filamentous Pezizomycotina fungi or other unassagined Ascomycota, some Basdidiomycota of the subphylum Agaricomycotina, and Muromycota. Although these ASVs are among the genera commonly detected in the human mycobiome, they often occur in a variety of other environments (e.g., *Exophiala*) or are typical saprophytes or phylopllans (e.g., *Aureobasidium*, *Penicillium*, *Cladosporium*, *Vishniacozyma*, *Mucor*) (Egbuta et al., 2016; Lebreton et al., 2020; A.-H. Li et al., 2020; Thitla et al., 2022). Some of these Pezizomycotina genera have been associated with the switch to a vegetarian diet in humans and are thought to be transient passengers ingested with food (Auchtung et al., 2018; Suhr et al., 2016). Furthermore, most of these subspecies variable ASVs also differed between transects, suggesting an environmental effect that was not detected by spatial analyses. Hypothetically, this could reflect patchily distributed microhabitats with different food resources that are unevenly represented in *musculus* and *domesticus* territories.

Although we identified differentially abundant bacterial ASVs responsible for the systematic community differences between the subspecies, and their biology pointed to transgenerational transmission, none of them showed signs of co-divergence with their hosts (i.e., the presence of closely related lineages, each restricted to one host subspecies). In contrast, they were all common to both subspecies, even when only the non-admixed hosts were considered. This may not be surprising, as the 16S rRNA marker used in this study may not have sufficient resolution to detect recently diverged microbes in the closely related host subspecies. Therefore, it is possible that these ASVs actually comprise co-diverging bacterial lineages and that the differences in their abundance reflect genetic differences between host subspecies. Consistent with the seminal work of (Falush et al., 2003) and (Linz et al., 2007) on human *Helicobacter* that co-diverged with human populations, a recent study by (Bendová et al., 2022) found two closely related *Helicobacter* ASVs with nearly perfect specificity for the two mouse subspecies under common garden conditions. However, our data from wild mice do not confirm the species specificity of these *Helicobacter* lineages. The *domesticus*-specific ASV of Bendová et al. (2022) was widespread in both subspecies, with a higher prevalence (47.2%) in *musclus* mice than in *domesticus* mice (27.6%), whereas the *musculus*-specific ASV was detected in only two *musculus* individuals at a single sampling site.

Body weight was a significant predictor of bacteriome and mycobiome composition and mycobiome diversity. Higher body mass was associated with increased abundance or prevalence of bacterial ASVs of the genera *Alistipes*, *Helicobacter*, *Odoribacter*, and unassigned Muribacualceae, whereas a negative association was observed only for the genus *Bacteroides* and the Lachnospiraceae NK4A136. This pattern is largely consistent with previous controlled experiments in which the abundance of *Alistipes* and *Odoribacter* was increased in mice fed with a high-fat diet (Walker et al., 2017), while *Bacteroidetes* and Lachnospiraceae NK4A136 were associated with a lean phenotype (Wu et al., 2020; Xiao et al., 2017). A positive effect on body weight was also observed with a single fungal ASV of the *Saccharomycotina* yeast from *Kazachstania*, the most dominant fungal genus in the entire data set. *Kazachstania* also dominated the mycobiomes of two species of free-living mice (*Mus musculus* and *M. spicilegus*) at a site approximately 600 km away (Bendová et al., 2020), suggesting that this genus may be a stable component of mouse mycobiomes. In humans, the most dominant fungi belong to the genus *Saccharomyces* (Nash et al., 2017), a close relative of *Kazachstania* (Shen et al., 2016). Interestingly, *Kazachastania* is present at very low levels or not at all in captive mice, while *Saccharomyces* is a common genus (Iliev et al., 2012; Mims et al., 2021) that has been strongly associated with metabolic disorders and weight gain (Mims et al., 2021). One possible explanation is that the mycobiome of captive mice can be humanized, as has been described for bacteriomes of captive animals (Clayton et al., 2016), and that the exchanged mycobiome members can confer the same functions to the host.

Contrary to laboratory mice (McGee & Huttenhower, 2021), we did not detect significant shifts in bacteriome composition between males and females, nor did we detect bacterial ASVs specifically associated with sex or pregnancy. However, in the mycobiome, the genera *Candida* and *Scopulariopsis* were more prevalent in females and *Aspergillus* was less common in pregnant females than in non-pregnant individuals.

Taken together, these results show that although the gut bacteriome of the wild house mouse depends on the host evolutionary history, host subspecies explain only a small part of the bacteriome variability. The effect of subspecies was even smaller for the gut mycobiome, but it is possible that the signal was blurred by high representation of dietary fungi. The existence of a major fraction of the microbiome that does not depend on the variables analysed is consistent with the hidden (i.e., below marker resolution) co-divergence of transgenerationally transmitted microbiota with their hosts, but also with the divergence of environmentally transmitted microbiota with the environment. In either case, selection pressures on microbiota composition and function appear to be uniform between host subspecies/different sites, as they are not reflected in the differential abundance of microbial lineages.

Host-associated microbiota can play a role in host post-copulatory reproductive isolation, if the microbiota in hybrids decreases host fitness relative to parental host lineages. Compelling evidence for this phenomenon comes from captive invertebrates (Brucker & Bordenstein, 2012, 2013). Theoretically, genes that specifically interact to produce a common fitness-related phenotype and diverge between allopatric host populations may be subject to Dobzhansky-Muller incompatibilities upon host admixture (Dobzhansky, 1937; Muller, 1942; Orr & Turelli, 2001). For example, if novel combinations of host genes that interact in regulating the composition and proper function of the host microbiota are incompatible in hybrids, the dysregulated microbiota will decrease hybrid fitness and suppress gene flow between parental host taxa. This mechanism has been invoked to explain significant shifts in bacterial communities in house mouse hybrids compared to similar communities in parental subspecies (J. Wang et al., 2015). To the incompatibilities between host genes, the hologenome theory of evolution (Brucker & Bordenstein, 2012, 2013; Zilber-Rosenberg & Rosenberg, 2008) also adds incompatibilities that may occur between specifically interacting, co-diverging genes of hosts and their microbes or of microbes alone. The possible presence of incompatibilities between mouse hosts and their microbiota stems from the study by Moeller et al. (2019), which showed negative health consequences of microbiomes exchanged between different mouse taxa.

Similar to the subspecies, the effect of admixture on gut microbiota was generally low in this study. Neither fungal nor bacterial alpha diversity changed with the degree of admixture, precluding the possibility that potential incompatibilities lead to depletion of the native microbiota or significant invasion of microbial species not normally present in non-admixed individuals. On the other hand, our data partially support the effect of admixture on microbial community composition, although this effect does not appear to be critical, in contrast to a previous report on house mice (J. Wang et al., 2015). Although whole community MDMR analyses revealed an effect of admixture only for the fungal community and the Jaccard dissimilarity matrix, we found a significant effect of admixture or the interaction of admixture with the subspecies for both the bacterial and fungal communities when focused on compositional variation along the major PCoA gradients of microbiota divergence. At the same time, the abundance or prevalence of some bacterial and fungal ASVs changed with admixture according to JSDM. In bacteriome, there were five ASVs whose prevalence was correlated with the degree of admixture, but the interaction between subspecies and admixture was not supported (Figure 4, Table S4). This means that the change in their prevalence with admixture had the same direction for both mouse subspecies, which is generally consistent with incompatibilities in the host genome. The admixed individuals were characterized by an increased prevalence of ASVs corresponding to *Alistipes*, *Bacteroides*, Lachnospiraceae, and *Helicobacter*, whereas another *Helicobacter* ASV was found less frequently in the admixed individuals. The interaction between admixture and subspecies, which could indicate incompatibilities between host and microbiota, was supported only for one bacterial ASV. This belonged to the genus *Lactobacillus*, and its prevalence showed a slightly negative association with admixture in *M. m. domesticus* (estimate = −0.1017, posterior support = 0.8140) and a positive association in *M. m. musculus* (0.4390, posterior support = 0.9960). Remarkably, we found no ASVs indicating pathological invasion of external bacteria in hybrids. On the contrary, all differentially prevalent bacteria were common house mouse commensals, which were also frequently detected in the non-admixed individuals.

A similar trend was observed for intestinal mycobiome, where the prevalence of four ASVs corresponding to *Penicillium* and unassigned Ascomycota and Basidiomycota decreased in admixed individuals of both mouse subspecies, whereas an *Exophiala* ASV showed the opposite pattern. In addition, the abundance or prevalence of five other fungal ASVs changed interactively with the admixture depending on the subspecies of their hosts.

Overall, the shifts in the microbial community and individual microbial lineages do not indicate strong dysregulation or pathogenic properties of the admixed microbiomes, suggesting that genetic incompatibilities, when present, have little impact on gut microbiomes in the two hybrid zone replicates studied.

In conclusion, our study demonstrated that failure to account for spatial autocorrelation, which is a common aspect of animal microbiota studies in free-living populations, can lead to misleading biological conclusions. Predictors of microbiota variation that were directly determined by the geographic distribution of collected samples often received higher statistical support when the spatial aspect was ignored in the analyses. After statistically controlling for spatial autocorrelation, the subspecies of mice and the extent of their genetic admixture had a significant effect on microbiota composition. However, the effect size of these two factors was rather small. Indeed, most of the differences between subspecies were due to gradual changes in the prevalence or abundance of a few microbial lineages common to both subspecies. Genetic admixture had only a minor effect on only several microbial ASVs (mostly on their prevalence but not on their abundance) that corresponded to common symbionts in free-living house mice. Also, at the whole community level, non-admixed and admixed individuals did not form separate clusters in the ordination analysis, in contrast to the previous report by (J. Wang et al., 2015). Overall, our data do not support the hypothesis that microbiota play an important role in host reproductive isolation in this particular model system.

## Acknowledgements

This study was supported by the Czech Science Foundation project no. 18–17796Y and 19-19307S. Computational resources were supplied by the project ‘e-Infrastruktura CZ’ (e-INFRA CZ LM2018140) supported by the Ministry of Education, Youth, and Sports of the Czech Republic. We are grateful to all our colleagues who participated in field sampling.

## Data Accessibility and Benefit-Sharing

The sequencing data associated with this project will be archived in the European Nucleotide Archive upon acceptance of the manuscript (project accession number: PRJEBXXXXX). The database of processed bacterial and fungal profiles, along with sample metadata, will be accessible through the repository https://github.com/jakubkreisinger. Benefits from this research accrue from the sharing of our data and results on public databases as described above.

## Author contributions

JK, DC, and JP designed the study; JP and MM secured field sampling and genotyping of mice; MK and BS provided extracted faecal DNA; LS and DC performed genotyping of the microbiota; JK performed data analysis; JK and DC drafted the manuscript; all authors edited the manuscript.

---

**Table S1:** Results of the alpha diversity analyses. Parameter estimates of the mixed models for the alpha diversity analyses. Variation in the two alpha diversity measures (Shannon index and Observed ASV richness) was assessed separately for bacterial and fungal communities. Models with and without spatial autocorrelation were evaluated in parallel. Predictors that were significant according to likelihood ratio tests are shown in bold.

| Community | Predictor | Response | Spatial model |  |  | Non-spatial model |  |  |
| --- | --- | --- | --- | --- | --- | --- | --- | --- |
|  |  |  | Estimate | SE | t.value | Estimate | SE | t.value |
| Bacteriome | (Intercept) | Observed | 1,8649 | 0,0501 | 37,2257 | 1,8649 | 0,0501 | 37,2257 |
|  | Subspecies (Domesticus vs Musculus) |  | -0,0239 | 0,0271 | -0,8827 | -0,0239 | 0,0271 | -0,8827 |
|  | Admixture |  | 0,0003 | 0,1013 | 0,0029 | 0,0003 | 0,1013 | 0,0029 |
|  | Subspecies (Domesticus vs Musculus) x Admix |  | 0,0344 | 0,1437 | 0,2393 | 0,0344 | 0,1437 | 0,2393 |
|  | Body weight |  | -0,0028 | 0,0024 | -1,1507 | -0,0028 | 0,0024 | -1,1507 |
|  | Sex+Gravidity (Females nongravid vs Females gravid) |  | -0,0041 | 0,0282 | -0,1466 | -0,0041 | 0,0282 | -0,1466 |
|  | Sex+Gravidity (Females nongravid vs Males) |  | -0,0247 | 0,0198 | -1,2483 | -0,0247 | 0,0198 | -1,2483 |
|  | Zone replicate (Czech vs Regensburg transect) |  | **-0,0835** | **0,0209** | **-3,9949** | **-0,0835** | **0,0209** | **-3,9949** |
|  | (Intercept) | Shannon | 3,1384 | 0,1469 | 21,3674 | 3,1384 | 0,1469 | 21,3679 |
|  | Subspecies (Domesticus vs Musculus) |  | 0,0788 | 0,0802 | 0,9823 | 0,0788 | 0,0802 | 0,9823 |
|  | Admixture |  | 0,4710 | 0,2990 | 1,5753 | 0,4710 | 0,2990 | 1,5753 |
|  | Subspecies (Domesticus vs Musculus) x Admix |  | -0,2882 | 0,4228 | -0,6817 | -0,2882 | 0,4228 | -0,6817 |
|  | Body weight |  | 0,0012 | 0,0070 | 0,1715 | 0,0012 | 0,0070 | 0,1715 |
|  | Sex+Gravidity (Females nongravid vs Females gravid) |  | 0,0323 | 0,0825 | 0,3919 | 0,0323 | 0,0825 | 0,3919 |
|  | Sex+Gravidity (Females nongravid vs Males) |  | 0,0039 | 0,0577 | 0,0680 | 0,0039 | 0,0577 | 0,0680 |
|  | Zone replicate (Czech vs Regensburg transect) |  | **-0,1422** | **0,0620** | **-2,2944** | **-0,1422** | **0,0620** | **-2,2945** |
| Mycobiome | (Intercept) | Observed | 1,1125 | 0,0822 | 13,5369 | 1,1100 | 0,0822 | 13,5043 |
|  | Subspecies (Domesticus vs Musculus) |  | 0,0491 | 0,0487 | 1,0077 | 0,0507 | 0,0485 | 1,0453 |
|  | Admixture |  | 0,0460 | 0,1821 | 0,2527 | 0,0423 | 0,1815 | 0,2330 |
|  | Subspecies (Domesticus vs Musculus) x Admix |  | -0,0231 | 0,2463 | -0,0937 | -0,0210 | 0,2459 | -0,0856 |
|  | Body weight |  | -0,0036 | 0,0039 | -0,9296 | -0,0036 | 0,0039 | -0,9204 |
|  | Sex+Gravidity (Females nongravid vs Females gravid) |  | 0,0549 | 0,0449 | 1,2235 | 0,0554 | 0,0450 | 1,2326 |
|  | Sex+Gravidity (Females nongravid vs Males) |  | -0,0233 | 0,0309 | -0,7517 | -0,0232 | 0,0310 | -0,7485 |
|  | Zone replicate (Czech vs Regensburg transect) |  | **-0,1579** | **0,0387** | **-4,0794** | **-0,1564** | **0,0385** | **-4,0668** |
|  | (Intercept) | Shannon | 1,6873 | 0,2109 | 7,9986 | 1,7357 | 0,2057 | 8,4377 |
|  | Subspecies (Domesticus vs Musculus) |  | **0,3426** | **0,1454** | **2,3559** | **0,3499** | **0,1318** | **2,6554** |
|  | Admixture |  | 0,8159 | 0,5176 | 1,5764 | 0,7850 | 0,4789 | 1,6391 |
|  | Subspecies (Domesticus vs Musculus) x Admix |  | -0,7698 | 0,6457 | -1,1922 | -0,6929 | 0,6344 | -1,0923 |
|  | Body weight |  | **-0,0253** | **0,0096** | **-2,6334** | **-0,0264** | **0,0096** | **-2,7383** |
|  | Sex+Gravidity (Females nongravid vs Females gravid) |  | 0,0878 | 0,1093 | 0,8031 | 0,0802 | 0,1100 | 0,7292 |
|  | Sex+Gravidity (Females nongravid vs Males) |  | -0,0614 | 0,0749 | -0,8196 | -0,0681 | 0,0751 | -0,9073 |
|  | Zone replicate (Czech vs Regensburg transect) |  | -0,0506 | 0,1274 | -0,3972 | -0,1007 | 0,1072 | -0,9398 |

**Table S2:** Variation in microbiota composition: major gradients approach. Likelihood ratio-based significance tests of the predictors of the mixed models for the two major PCoA gradients in microbiota composition. Significance (p) was derived from deviance changes (χ^2^) and corresponding degrees of freedom (Δ D.f.). Variation in scores for the first two PCoA axes for two beta diversity measures (Bray-Curtis and Jaccard) was assessed separately for bacterial and fungal profiles. Models with and without spatial autocorrelation were fitted in parallel. Values of significant predictors are in bold.

| Community | Dissimilarity measure | PcoA axis | Predictor | $\Delta$ D.f. | Spatial models | | Non-spatial models | |
| --- | --- | --- | --- | --- | --- | --- | --- | --- |
| | | | | | $\chi^2$ | p | $\chi^2$ | p |
| Bacteriome | Bray-Curtis | axis1 | Subspecies | 1 | **12,9609** | **0,0003** | **32,8071** | **0,0000** |
|  |  |  | Admixture | 1 | 0,4698 | 0,4931 | 1,3619 | 0,2432 |
|  |  |  | Subspec x Admix | 1 | 1,1033 | 0,2935 | 1,4078 | 0,2354 |
|  |  |  | Body weight | 1 | **4,9435** | **0,0262** | **4,4330** | **0,0353** |
|  |  |  | Sex+Gravidity | 2 | 1,4191 | 0,4919 | 1,1393 | 0,5657 |
|  |  |  | Zone replicate | 1 | **4,3203** | **0,0377** | **15,9884** | **0,0001** |
|  |  | axis2 | Subspecies | 1 | **4,4313** | **0,0353** | **10,3771** | **0,0013** |
|  |  |  | Admixture | 1 | **1,5148** | **0,2184** | **5,4107** | **0,0200** |
|  |  |  | Subspec x Admix | 1 | **5,9027** | **0,0151** | **5,1397** | **0,0234** |
|  |  |  | Body weight | 1 | 0,4163 | 0,5188 | 0,7922 | 0,3734 |
|  |  |  | Sex+Gravidity | 2 | 0,4622 | 0,7937 | 0,6412 | 0,7257 |
|  |  |  | Zone replicate | 1 | **6,2561** | **0,0124** | **5,0363** | **0,0248** |
|  | Jaccard | axis1 | Subspecies | 1 | **13,2820** | **0,0003** | **29,2833** | **0,0000** |
|  |  |  | Admixture | 1 | 0,3218 | 0,5705 | 1,1915 | 0,2750 |
|  |  |  | Subspec x Admix | 1 | 0,5528 | 0,4572 | 0,5956 | 0,4402 |
|  |  |  | Body weight | 1 | **4,1735** | **0,0411** | 3,7883 | 0,0516 |
|  |  |  | Sex+Gravidity | 2 | 1,7538 | 0,4161 | 1,4517 | 0,4839 |
|  |  |  | Zone replicate | 1 | **13,2820** | **0,0003** | **16,8027** | **0,0000** |
|  |  | axis2 | Subspecies | 1 | **4,4430** | **0,0350** | 0,8152 | 0,3666 |
|  |  |  | Admixture | 1 | 0,9066 | 0,3410 | **4,5587** | **0,0328** |
|  |  |  | Subspec x Admix | 1 | 1,2822 | 0,2575 | **4,1233** | **0,0423** |
|  |  |  | Body weight | 1 | 0,7977 | 0,3718 | 1,1463 | 0,5637 |
|  |  |  | Sex+Gravidity | 2 | 1,0362 | 0,5957 | 1,1329 | 0,2872 |
|  |  |  | Zone replicate | 1 | 3,6489 | 0,0561 | 0,9954 | 0,3184 |
| Mycobiome | Bray-Curtis | axis1 | Subspecies | 1 | **0,6241** | **0,4295** | **1,3578** | **0,2439** |
|  |  |  | Admixture | 1 | **0,0032** | **0,9549** | **0,1793** | **0,6720** |
|  |  |  | Admix x Subspec | 1 | **4,6292** | **0,0314** | **5,9696** | **0,0146** |
|  |  |  | Body weight | 1 | 0,2109 | 0,6461 | 0,4719 | 0,4921 |
|  |  |  | Sex+Gravidity | 2 | 1,0960 | 0,5781 | 1,0321 | 0,5969 |
|  |  |  | Zone replicate | 1 | **13,5821** | **0,0002** | **31,4592** | **0,0000** |
|  |  | axis2 | Subspecies | 1 | 2,9698 | 0,0848 | **9,2542** | **0,0023** |
|  |  |  | Admixture | 1 | 2,8420 | 0,0918 | 1,2092 | 0,2715 |
|  |  |  | Admix x Subspec | 1 | 0,5317 | 0,4659 | 1,0248 | 0,3114 |
|  |  |  | Body weight | 1 | **8,4946** | **0,0036** | 0,9329 | 0,3341 |
|  |  |  | Sex+Gravidity | 2 | 0,1205 | 0,9415 | 0,0877 | 0,9571 |
|  |  |  | Zone replicate | 1 | 1,0990 | 0,2945 | 0,9329 | 0,3341 |
|  | Jaccard | axis1 | Subspecies | 1 | 0,0356 | 0,8503 | 0,0026 | 0,9596 |
|  |  |  | Admixture | 1 | **8,8342** | **0,0030** | **8,4839** | **0,0036** |
|  |  |  | Admix x Subspec | 1 | **4,3497** | **0,0370** | **4,7429** | **0,0294** |
|  |  |  | Body weight | 1 | 0,0053 | 0,9418 | 0,0356 | 0,8503 |
|  |  |  | Sex+Gravidity | 2 | 2,0669 | 0,3558 | 2,0366 | 0,3612 |
|  |  |  | Zone replicate | 1 | 1,4695 | 0,2254 | 2,4980 | 0,1140 |
|  |  | axis2 | Subspecies | 1 | 0,1798 | 0,6716 | 0,6576 | 0,4174 |
|  |  |  | Admixture | 1 | 1,2715 | 0,2595 | 2,5847 | 0,1079 |
|  |  |  | Admix x Subspec | 1 | 0,8324 | 0,3616 | 0,8018 | 0,3706 |
|  |  |  | Body weight | 1 | 1,1934 | 0,2746 | 2,8659 | 0,0905 |
|  |  |  | Sex+Gravidity | 2 | **8,2974** | **0,0158** | **6,4049** | **0,0407** |
|  |  |  | Zone replicate | 1 | **34,4841** | **0,0000** | **8,4839** | **0,0036** |

**Table S3:**
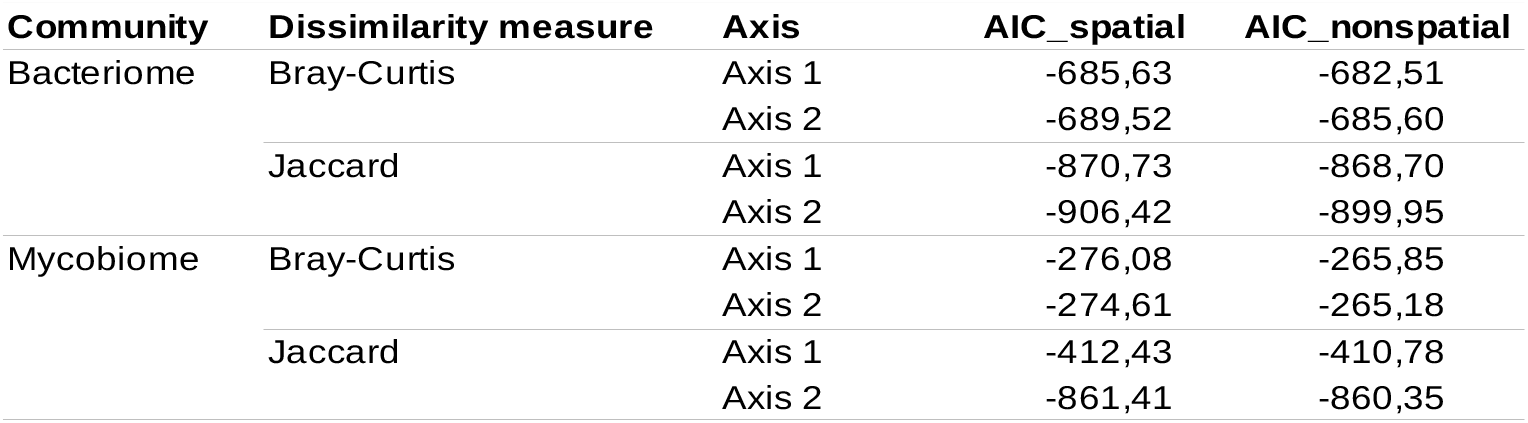
Comparison of spatial and non-spatial mixed models of microbiota composition. Mixed models with scores for the first two PCoA axes calculated based on Bray-Curtis and Jaccard dissimilarities and controlled for spatial autocorrelation were compared with their simplified counterparts in which spatial distances between samples were ignored. The comparison was based on the AIC values for each model.

| Community | Dissimilarity measure | Axis | AIC_spatial | AIC_nonspatial |
| --- | --- | --- | --- | --- |
| Bacteriome | Bray-Curtis | Axis 1 | -685,63 | -682,51 |
|  |  | Axis 2 | -689,52 | -685,60 |
|  | Jaccard | Axis 1 | -870,73 | -868,70 |
|  |  | Axis 2 | -906,42 | -899,95 |
| Mycobiome | Bray-Curtis | Axis 1 | -276,08 | -265,85 |
|  |  | Axis 2 | -274,61 | -265,18 |
|  | Jaccard | Axis 1 | -412,43 | -410,78 |
|  |  | Axis 2 | -861,41 | -860,35 |

**Table S4:** Parameters of the mixed models of microbiota composition from the major gradients approach. Variation in scores for the first two PCoA axes for two beta diversity measures (Bray-Curtis and Jaccard) was assessed separately for bacterial and fungal profiles. Models with and without spatial autocorrelation were evaluated in parallel.

| Community | Dissimilarity | PcoA axis | Predictor | Spatial models |  |  | Non-spatial models |  |  |
| --- | --- | --- | --- | --- | --- | --- | --- | --- | --- |
|  |  |  |  | Estimate | Cond. SE | t.value | Estimate | Cond. SE | t.value |
| Bacteriome | Bray-Curtis | axis1 | (Intercept) | 0,0885 | 0,0339 | 2,6078 | 0,0800 | 0,0320 | 2,5049 |
|  |  |  | Subspecies (Domesticus vs Musculus) | **-0,1029** | **0,0243** | **-4,2379** | **-0,1106** | **0,0203** | **-5,4610** |
|  |  |  | Admixture | -0,0915 | 0,0788 | -1,1605 | -0,1181 | 0,0719 | -1,6440 |
|  |  |  | Subspecies (Domesticus vs Musculus) x Admix | 0,1085 | 0,1016 | 1,0680 | 0,1163 | 0,0974 | 1,1933 |
|  |  |  | Body weight | **-0,0030** | **0,0015** | **-2,0530** | **-0,0029** | **0,0015** | **-1,9509** |
|  |  |  | Sex+Gravidity (Females nongravid vs Females gravid) | -0,0115 | 0,0170 | -0,6755 | -0,0107 | 0,0171 | -0,6276 |
|  |  |  | Sex+Gravidity (Females nongravid vs Males) | -0,0133 | 0,0117 | -1,1381 | -0,0119 | 0,0117 | -1,0175 |
|  |  |  | Zone replicate (Czech vs Regensburg transect) | **0,0626** | **0,0222** | **2,8178** | **0,0738** | **0,0163** | **4,5325** |
|  |  | axis2 | (Intercept) | 0,0429 | 0,0314 | 1,3665 | 0,0342 | 0,0309 | 1,1082 |
|  |  |  | Subspecies (Domesticus vs Musculus) | **-0,0665** | **0,0200** | **-3,3244** | **-0,0699** | **0,0183** | **-3,8212** |
|  |  |  | Admixture | -0,1965 | 0,0709 | -2,7689 | -0,2258 | 0,0666 | -3,3916 |
|  |  |  | Subspecies (Domesticus vs Musculus) x Admix | **0,2271** | **0,0924** | **2,4570** | **0,2108** | **0,0919** | **2,2942** |
|  |  |  | Body weight | 0,0009 | 0,0014 | 0,6163 | 0,0012 | 0,0015 | 0,8471 |
|  |  |  | Sex+Gravidity (Females nongravid vs Females gravid) | 0,0028 | 0,0167 | 0,1678 | 0,0035 | 0,0169 | 0,2083 |
|  |  |  | Sex+Gravidity (Females nongravid vs Males) | 0,0085 | 0,0116 | 0,7307 | 0,0098 | 0,0117 | 0,8430 |
|  |  |  | Zone replicate (Czech vs Regensburg transect) | **-0,0390** | **0,0171** | **-2,2845** | **-0,0259** | **0,0144** | **-1,7970** |
|  | Jaccard | axis1 | (Intercept) | 0,0642 | 0,0270 | 2,3755 | 0,0588 | 0,0259 | 2,2721 |
|  |  |  | Subspecies (Domesticus vs Musculus) | **-0,0744** | **0,0184** | **-4,0344** | **-0,0782** | **0,0160** | **-4,8783** |
|  |  |  | Admixture | -0,0580 | 0,0618 | -0,9388 | -0,0757 | 0,0574 | -1,3187 |
|  |  |  | Subspecies (Domesticus vs Musculus) x Admix | 0,0607 | 0,0809 | 0,7500 | 0,0607 | 0,0783 | 0,7753 |
|  |  |  | Body weight | **-0,0024** | **0,0012** | **-1,9473** | -0,0023 | 0,0012 | -1,8637 |
|  |  |  | Sex+Gravidity (Females nongravid vs Females gravid) | -0,0071 | 0,0139 | -0,5119 | -0,0067 | 0,0140 | -0,4780 |
|  |  |  | Sex+Gravidity (Females nongravid vs Males) | -0,0122 | 0,0096 | -1,2742 | -0,0115 | 0,0096 | -1,1922 |
|  |  |  | Zone replicate (Czech vs Regensburg transect) | **0,0515** | **0,0164** | **3,1344** | **0,0586** | **0,0128** | **4,5820** |
|  |  | axis2 | (Intercept) | 0,0010 | 0,0279 | 0,0359 | 0,0082 | 0,0246 | 0,3343 |
|  |  |  | Subspecies (Domesticus vs Musculus) | 0,0139 | 0,0148 | 0,9442 | 0,0210 | 0,0107 | 1,9709 |
|  |  |  | Admixture | -0,1005 | 0,0881 | -1,1415 | 0,0273 | 0,0618 | 0,4421 |
|  |  |  | Subspecies (Domesticus vs Musculus) x Admix | 0,2152 | 0,1079 | 1,9954 | 0,0752 | 0,0823 | 0,9137 |
|  |  |  | Body weight | -0,0008 | 0,0011 | -0,7281 | -0,0012 | 0,0012 | -1,0608 |
|  |  |  | Sex+Gravidity (Females nongravid vs Females gravid) | -0,0045 | 0,0133 | -0,3419 | -0,0047 | 0,0134 | -0,3483 |
|  |  |  | Sex+Gravidity (Females nongravid vs Males) | -0,0084 | 0,0092 | -0,9084 | -0,0098 | 0,0093 | -1,0539 |
|  |  |  | Zone replicate (Czech vs Regensburg transect) | 0,0014 | 0,0239 | 0,0566 | -0,0023 | 0,0162 | -0,1443 |

**Table S4 – Mycobiome**
| Community | Dissimilarity | PcoA axis | Predictor | Spatial models |  |  | Non-spatial models |  |  |
| --- | --- | --- | --- | --- | --- | --- | --- | --- | --- |
|  |  |  |  | Estimate | Cond. SE | t.value | Estimate | Cond. SE | t.value |
| Mycobiome | Bray-Curtis | axis1 | (Intercept) | 0,1135 | 0,0558 | 2,0346 | 0,1017 | 0,0540 | 1,8832 |
|  |  |  | Subspecies (Domesticus vs Musculus) | -0,0942 | 0,0452 | -2,0817 | -0,0993 | 0,0401 | -2,4740 |
|  |  |  | Admixture | -0,1914 | 0,1438 | -1,3317 | -0,2613 | 0,1357 | -1,9256 |
|  |  |  | Subspecies (Domesticus vs Musculus) x Admix | **0,3829** | **0,1732** | **2,2108** | **0,4321** | **0,1729** | **2,4986** |
|  |  |  | Body weight | 0,0010 | 0,0024 | 0,4280 | 0,0016 | 0,0024 | 0,6788 |
|  |  |  | Sex+Gravidity (Females nongravid vs Females gravid) | 0,0111 | 0,0270 | 0,4125 | 0,0075 | 0,0275 | 0,2736 |
|  |  |  | Sex+Gravidity (Females nongravid vs Males) | 0,0192 | 0,0183 | 1,0480 | 0,0185 | 0,0185 | 1,0002 |
|  |  |  | Zone replicate (Czech vs Regensburg transect) | **-0,1842** | **0,0407** | **-4,5197** | **-0,1746** | **0,0341** | **-5,1171** |
|  |  | axis2 | (Intercept) | -0,0336 | 0,0565 | -0,5942 | -0,0358 | 0,0520 | -0,6885 |
|  |  |  | Subspecies (Domesticus vs Musculus) | -0,0839 | 0,0445 | -1,8871 | **-0,0936** | **0,0346** | **-2,7089** |
|  |  |  | Admixture | -0,2616 | 0,1428 | -1,8314 | -0,1862 | 0,1237 | -1,5053 |
|  |  |  | Subspecies (Domesticus vs Musculus) x Admix | 0,1319 | 0,1720 | 0,7671 | 0,1652 | 0,1622 | 1,0182 |
|  |  |  | Body weight | **0,0062** | **0,0024** | **2,6071** | 0,0062 | 0,0024 | 2,5600 |
|  |  |  | Sex+Gravidity (Females nongravid vs Females gravid) | 0,0090 | 0,0271 | 0,3304 | 0,0099 | 0,0275 | 0,3610 |
|  |  |  | Sex+Gravidity (Females nongravid vs Males) | 0,0055 | 0,0185 | 0,2962 | 0,0030 | 0,0187 | 0,1590 |
|  |  |  | Zone replicate (Czech vs Regensburg transect) | -0,0146 | 0,0417 | -0,3495 | -0,0145 | 0,0284 | -0,5094 |
|  | Jaccard | axis1 | (Intercept) | -0,0502 | 0,0421 | -1,1929 | -0,0502 | 0,0421 | -1,1929 |
|  |  |  | Subspecies (Domesticus vs Musculus) | 0,0423 | 0,0250 | 1,6951 | 0,0423 | 0,0250 | 1,6951 |
|  |  |  | Admixture | **0,3633** | **0,0933** | **3,8931** | **0,3633** | **0,0933** | **3,8931** |
|  |  |  | Subspecies (Domesticus vs Musculus) x Admix | **-0,2963** | **0,1262** | **-2,3479** | **-0,2963** | **0,1262** | **-2,3479** |
|  |  |  | Body weight | 0,0004 | 0,0020 | 0,1914 | 0,0004 | 0,0020 | 0,1914 |
|  |  |  | Sex+Gravidity (Females nongravid vs Females gravid) | -0,0280 | 0,0230 | -1,2151 | -0,0280 | 0,0230 | -1,2151 |
|  |  |  | Sex+Gravidity (Females nongravid vs Males) | -0,0196 | 0,0158 | -1,2357 | -0,0196 | 0,0158 | -1,2357 |
|  |  |  | Zone replicate (Czech vs Regensburg transect) | 0,0263 | 0,0198 | 1,3273 | 0,0263 | 0,0198 | 1,3273 |
|  |  | axis2 | (Intercept) | 0,0339 | 0,0259 | 1,3076 | 0,0314 | 0,0253 | 1,2409 |
|  |  |  | Subspecies (Domesticus vs Musculus) | 0,0177 | 0,0167 | 1,0609 | 0,0193 | 0,0151 | 1,2775 |
|  |  |  | Admixture | 0,0912 | 0,0611 | 1,4928 | 0,0953 | 0,0564 | 1,6900 |
|  |  |  | Subspecies (Domesticus vs Musculus) x Admix | -0,0709 | 0,0773 | -0,9172 | -0,0684 | 0,0761 | -0,8984 |
|  |  |  | Body weight | 0,0014 | 0,0012 | 1,1550 | 0,0013 | 0,0012 | 1,0650 |
|  |  |  | Sex+Gravidity (Females nongravid vs Females gravid) | **0,0326** | **0,0137** | **2,3792** | **0,0336** | **0,0138** | **2,4378** |
|  |  |  | Sex+Gravidity (Females nongravid vs Males) | **0,0123** | **0,0095** | **1,2957** | **0,0122** | **0,0095** | **1,2875** |
|  |  |  | Zone replicate (Czech vs Regensburg transect) | **-0,2073** | **0,0146** | **-14,2106** | **-0,2061** | **0,0120** | **-17,1034** |

**Table S5:**
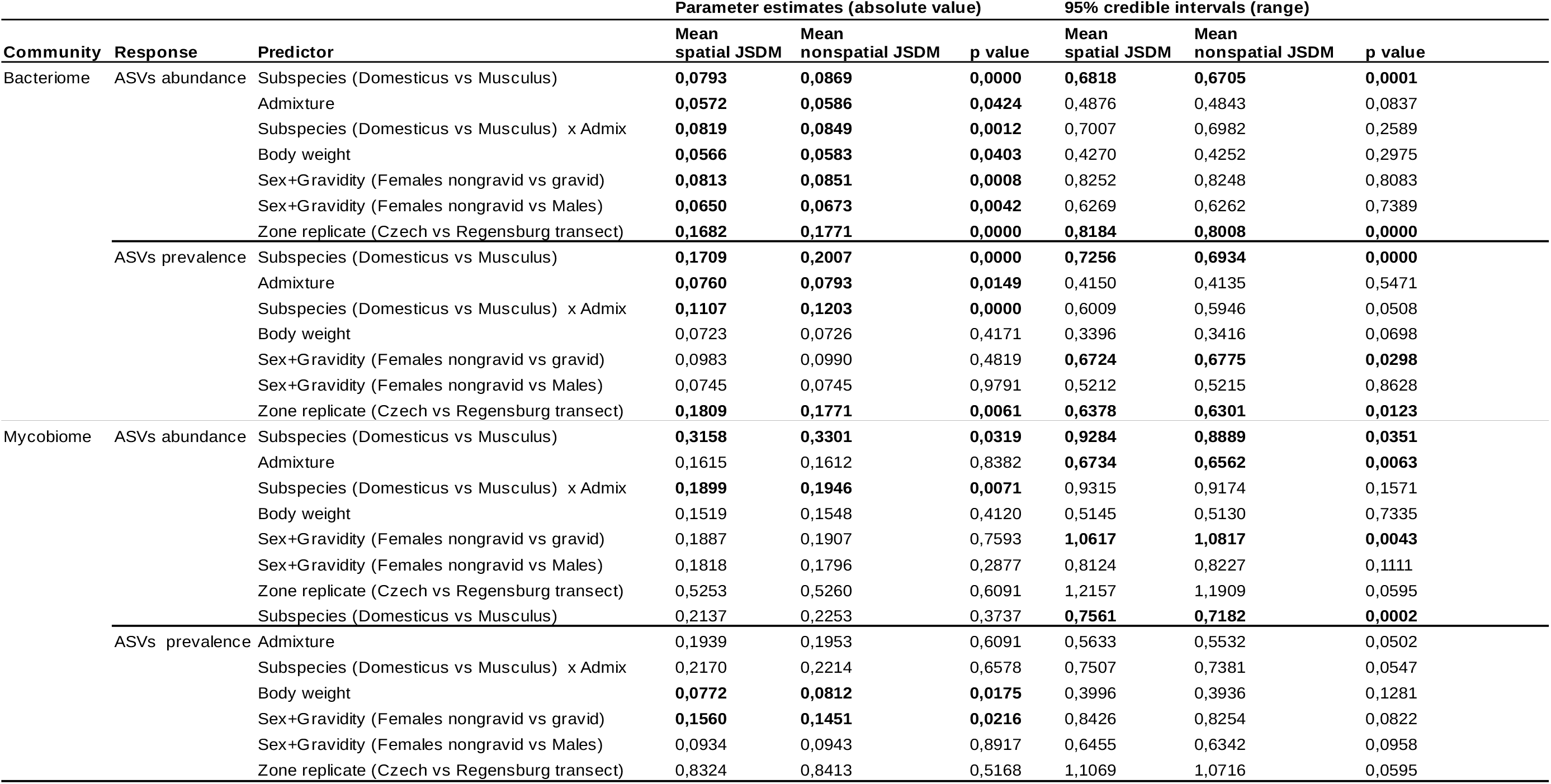
Comparison of parameter estimates and their uncertainties for spatial and non-spatial versions of the joint species distribution models (JSDM) For each predictor, all ASVs included in the model were considered in the comparisons. For the spatial and non-spatial JSDMs, absolute values of parameter estimates and their uncertainties, expressed as 95% posterior credible intervals, were compared using the Wilcoxon paired test. Significant differences (p < 0.05) are shown in bold.

**Figure S1:**
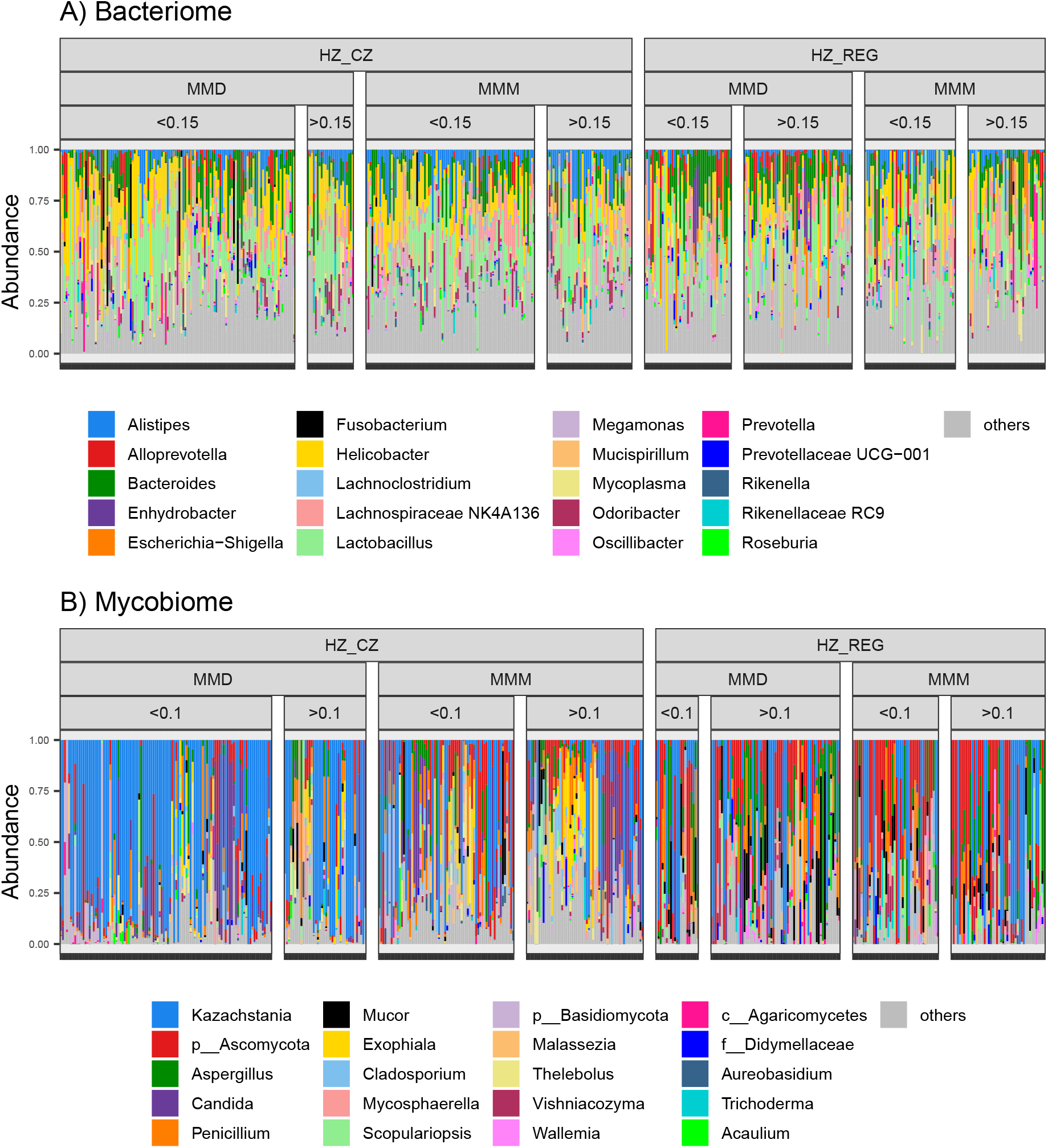
Taxonomic composition of the gut bacteriome and mycobiome at the genus level. Proportion of dominant A) bacterial and B) fungal genera in two house mouse subspecies (MMD = M. m. domesticus; MMM = M. m. musculus) and in two hybrid zone replicates (HZ_CZ = Czech transect; HZ_REG = Regensburg transect). For this plot, mice were ranked by degree of genetic admixture (< 0.1; > 0.1), but for analyses, degree of admixture was used as a continuous variable.

**Figure S2:**
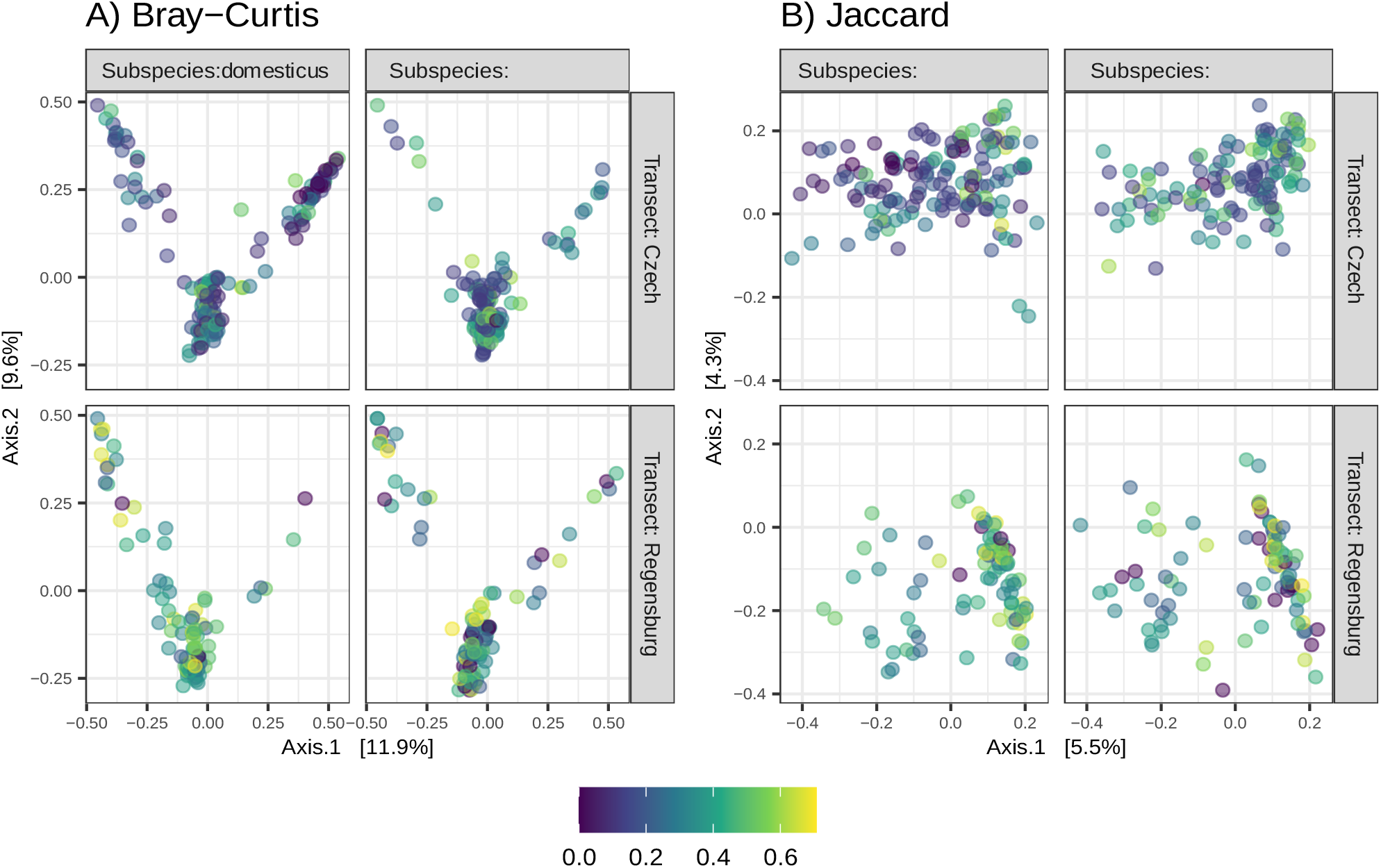
Ordination of gut mycobiome. PCoA ordination showing GM divergence among hybrid zone relicates (Transect: Czech or Regensburg), house mouse subspecies (Subspecies: domesticus or musculus), and due to genetic admixture (color scale express degree of admixture after square root transformation, suppressing the influence of extreme values). Ordination was performed for dissimilarities between samples accounting for A) variation in relative ASV frequencies (Bray-Curtis) and B) presence/absence of ASVs (Jaccard).

**Figure S3:**
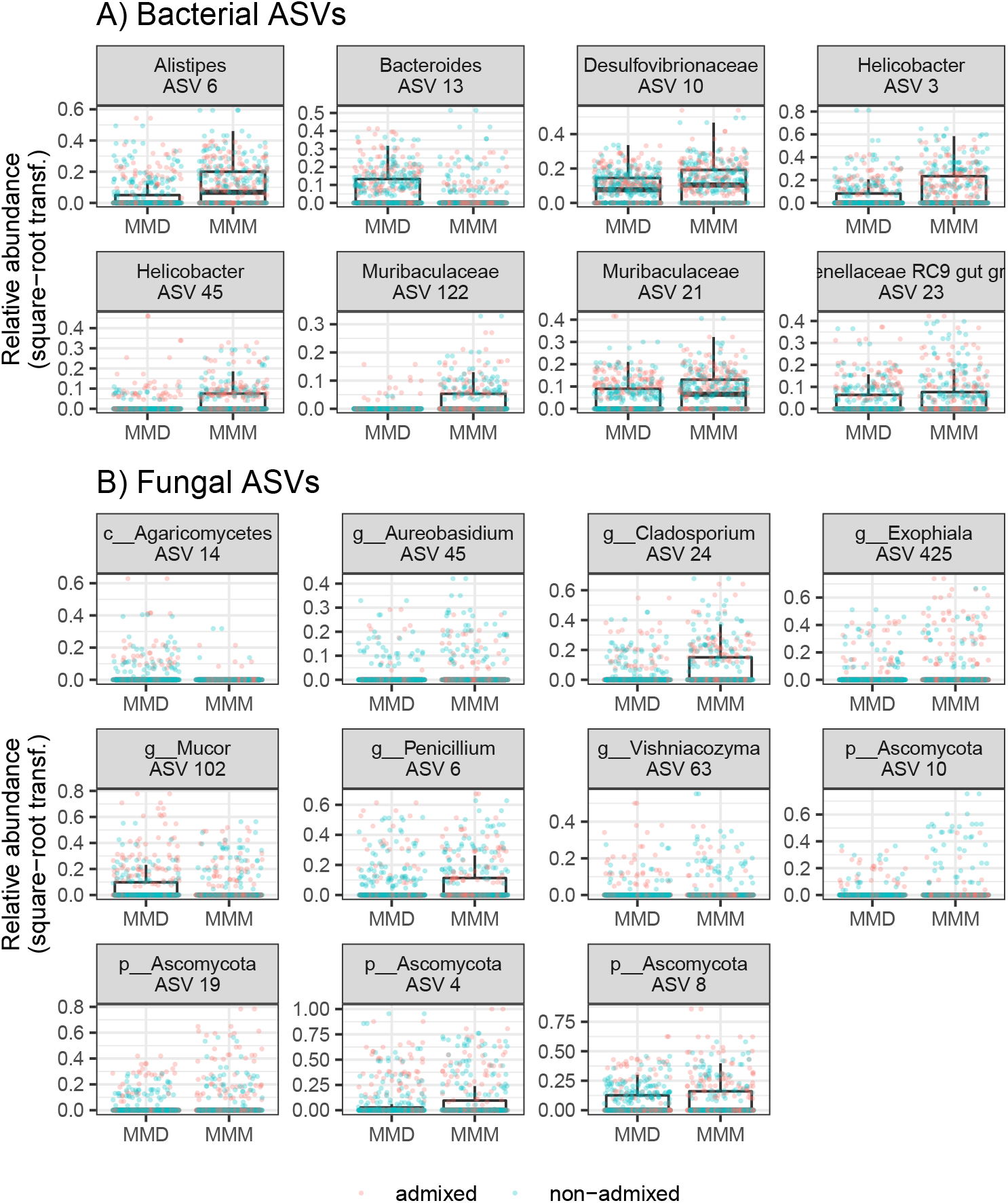
ASVs that differ between house mouse subspecies. Relative abundance of A) bacterial B) fungal ASVs showing differential abundance between *M. m. musculus* (MMM) and *M. m. domesticus* (MMD) according to JSDM. Colours separate individuals with high (> 0.1 = admixed) and low (< 0.1 = non-admixed) levels of genetic admixture. The admixture was divided into the two categories for display purposes only.

